# RNA-targeted aminopyridine compounds provide a new antifungal strategy

**DOI:** 10.64898/2026.11.04.764270

**Authors:** Jimin Hwang, Luke J. Sisto, Tao Shi, Michael C. Van Zandt, Anna Marie Pyle

## Abstract

While RNA is an emerging therapeutic target class, progress is limited by a shortage of well-defined targets in which ligand binding is coupled to measurable function. The group I intron, a self-splicing ribozyme essential in fungal mitochondria but absent from humans, offers both a tractable platform for exploring first principles of small molecule-RNA targeting and a selective antifungal target. We recalibrated an intron-splicing assay to approximate cellular conditions and performed an extensive, cryo-EM-guided structure-activity relationship campaign against the *Candida albicans* mtLSU intron. Ligands containing a flexible diamine outperformed conformationally constrained analogues, suggesting a general principle for engaging dynamic RNA surfaces and yielding lead aminopyridine **56** (IC□□21.6 nM). Compound **56** synergizes with amphotericin B against *C. albicans*, suggesting a path for combination therapies. RT-qPCR confirmed direct target engagement in cells. This work establishes the fungal group I intron as a viable, mechanistically distinct antifungal target and a model platform for RNA-targeted discovery.

## Introduction

RNA has emerged as a valuable new target class in drug discovery. Although most FDA-approved small molecule drugs interact with proteins, protein-coding sequences account only for ∼2% of the human genome, and only a fraction of the encoded proteome is considered druggable.^1,2^ The remainder of the transcriptome is composed of non-coding RNAs or segments of RNA with regulatory functions, many of which are linked to disease or oncogenesis. In some cases, the gene of interest does not encode a protein product suitable for drug targeting, making it essential to focus on the RNA itself.^2,3^ Messenger RNAs represent attractive targets because they contain a variety of structured motifs in UTRs, introns and even ORFs. Furthermore, a single mRNA transcript yields 10³–10□ protein copies, making inhibition of mRNA function a more efficient way to modulate gene expression than engagement with downstream protein products.^4,5^

RNA targeting modalities have expanded from complementary oligonucleotides that target primary sequence to small molecules, which recognize specific RNA structures.^6^ RNA molecules adopt defined secondary and tertiary structures, both locally and globally.^7,8^ Although some of these structures are conformationally dynamic, in RNA, these dynamics can reflect interconversion among well-defined folds rather than a lack of well-formed structure. Locally stable motifs can provide discrete and ligand-accessible pockets, or even cryptic pockets that become stable upon ligand binding.^9^ RNA motifs containing specific structures can selectively recognize small molecules with high affinity and a growing number of RNAs have now been validated as small molecule targets.^10–18^ Databases such as R-BIND have helped to define the chemical characteristics common to RNA-targeting small molecules.^19,20^ However, because available RNA recognition data derive from a limited number of biochemically or structurally-characterized RNA targets (most of them relatively small), expanding both the range of targetable RNA classes and the chemical space of their binders remain important goals.^21^ Identifying systems that provide foundational information on RNA-small molecule recognition is crucial, as productive targeting can be achieved and rationalized when RNA presents both a structurally defined pocket and a clear functional understanding of the features that drive ligand binding.

Certain types of self-splicing introns satisfy these criteria by combining well-characterized structures with clear functional outputs.^14,22,23^ The group I intron is a self-splicing ribozyme that excises itself through two transesterification reactions that are initiated by an exogenous guanosine cofactor that binds within a highly conserved active-site.^24,25^ Subsequent catalysis is strictly dependent on correct folding of the intron RNA architecture surrounding the guanosine binding site. A ribozyme in a misfolded, kinetically trapped conformation cannot react, so conformational state is reported via the overall catalytic activity.^26,27^ Group I introns are well suited to inhibitor discovery because ligands that shift the intron conformation produce a measurable change in splicing activity, resulting in a direct functional readout that is rarely available for other RNA targets. We recently established that a group I intron found within the *Candida albicans* mitochondrial large subunit gene (C.a.mtLSU), is readily druggable and we conducted preliminary SAR to develop an aminopyrimidine scaffold.^23^ Importantly, that study was accompanied by high resolution ligand-bound structures that laid the foundation for subsequent stages of SAR and rational optimization.

Beyond their value as informative RNA-targeting platforms, group I introns represent attractive therapeutic targets. Group I intron splicing is essential for RNA processing in fungal mitochondrial genomes, including those of major fungal pathogens.^15^ Because group I introns are selective and absent from animal genomes, these introns represent particularly valuable drug targets.^22,28–30^ Invasive fungal infections represent an increasing threat to public health, yet currently available antifungal drugs act by disrupting fungal cell walls or membrane integrity.

Resistance arises rapidly because these agents converge on a narrow set of targets within the same two pathways.^31,32^ By contrast, group I intron splicing is required for expression of mitochondrially encoded respiratory-chain components, via an active site that is conserved and mutationally constrained. Therefore, group I intron inhibition acts through a mechanism distinct from that of current antifungals and which may avoid rapid development of resistance. Moreover, by impairing mitochondrial bioenergetics, inhibition of group I introns compromises ATP-dependent efflux pumps that drive resistance, enabling group I intron-targeted drugs to act synergistically with existing antifungals.

During the course of our previous work, we identified an aminopyrimidine compound class that binds the intron active site and stabilizes a non-catalytic conformation of C.a.mtLSU intron.^23^ While that work laid a valuable foundation, SAR was limited and did not leverage high resolution structural information for downstream optimization of the compound family. Second, the HTS-compatible assay was conducted under conditions that do not reflect physiologically optimal RNA folding or guanosine cofactor conditions. Perhaps most importantly, previous aminopyrimidine compounds were not evaluated as antifungal candidates, leaving the group I intron unvalidated as a drug target. In this study, we have addressed these issues and expanded our knowledge of intron targeting by re-engineering the splicing assay to test inhibitors against a pre-folded, catalytically competent ribozyme under physiological cofactor conditions. We then conducted a structure-guided SAR campaign to characterize small molecule recognition at the active site. Finally, we evaluated the optimized compounds in fungal growth assays, establishing the group I intron as a viable antifungal target.

## RESULTS

### Optimization of the C.a.mtLSU intron splicing inhibition assay

To establish biochemical conditions that better approximate the intracellular environment, we optimized the RNA folding protocol, cofactor identity and concentrations for the C.a.mtLSU intron splicing assay. Previous work utilized rapid cooling following denaturation, a procedure that can trap RNAs in misfolded states, as well-documented for other group I introns.^33^ Moreover, the inhibitor had been added simultaneously with DMSO during a DMSO-mediated refolding step, exposing the small molecule to the RNA before it adopted a catalytically competent conformation. To address these issues before developing the compounds further, we re-optimized the folding step to include a slow-cooling ramp, ensuring the RNA adopts a catalytically active conformation prior to adding DMSO or small molecule inhibitors (Supplementary Fig. S1).

We next addressed cofactor identities and concentrations. Acting as competitive inhibitors, aminopyrimidine compounds compete with the guanosine cofactor in group I introns, making it important to utilize the form of guanosine that is typically present in the cell.^23,34^ Previous studies used 20 µM uncharged guanosine as the splicing cofactor, which enhanced the sensitivity of initial screening. However, these conditions do not reflect the intracellular environment, where GTP—the predominant guanosine component in cells—is present at concentrations of 0.1 to 1 mM. Because GTP and guanosine differ in their ability to facilitate group I intron splicing, their competition with small molecule inhibitors is also expected to be different. Given that we would ultimately aim to deploy aminopyrimidines as antifungals, we therefore redesigned the splicing assay using GTP.

We first compared C.a.mtLSU intron splicing kinetics under guanosine- and GTP-based conditions at the same concentration (Fig. 1C). At 20 µM guanosine, the precursor conversion rate constant (*k*_obs_) was 0.071 min^-1^, whereas 20 µM GTP gave slower splicing with a *k*_obs_ of 0.029 min^-1^. Increasing GTP to 200 µM produced monophasic splicing with a *k*_obs_ of 0.14 min^-1^. Although GTP exhibits a lower catalytic efficiency than guanosine at equivalent concentrations, the C.a.mtLSU intron retains robust kinetic efficiency in GTP and reaches near-maximal terminal yields under physiologically relevant GTP concentrations (Fig. 1C).

**Figure 1.**
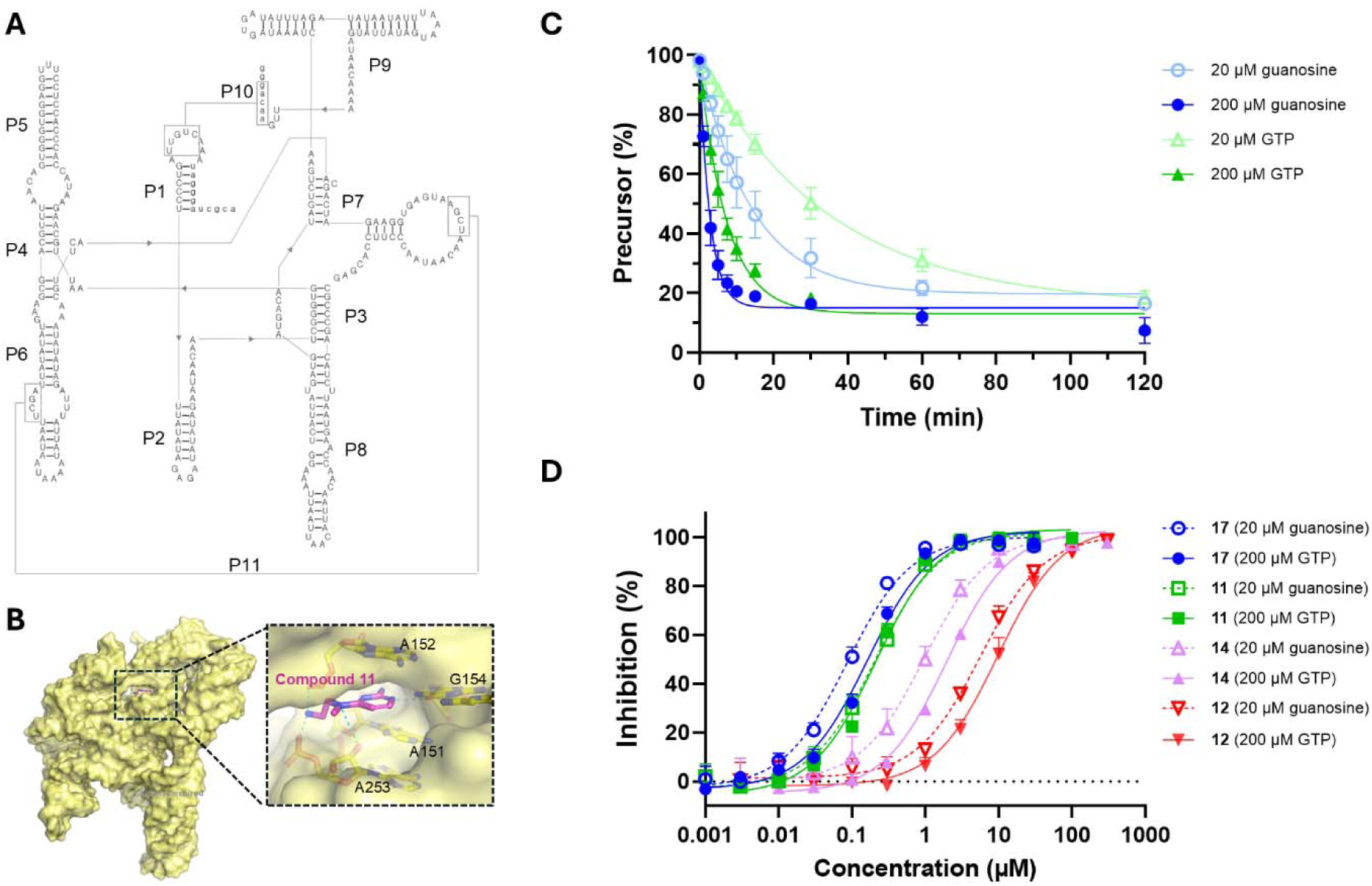
C.a.mtLSU group I intron-targeting aminopyrimidines inhibit intron-mediated splicing under physiologically relevant conditions. (A) Secondary structure of the C.a.mtLSU intron, the therapeutic target for antifungal agent development. (B) Cryo-EM structure of an aminopyrimidine, compound 11, bound to the guanosine-binding pocket of C.a.mtLSU (PDB ID: 9MQT). (C) Optimization of *in vitro* splicing conditions to approximate intracellular nucleotide environments. *k*_obs_ of 20 μM guanosine, 200 μM guanosine, 20 μM GTP, 200 μM GTP is 0.071, 0.35, 0.029, 0.14 min^-1^ respectively. (D) Dose–response curves for aminopyrimidine-mediated splicing inhibition under previously reported conditions (dotted lines) and optimized conditions in this study (solid lines)

To compare the trends in structure-activity relationship (SAR) between the 20 µM guanosine and 200 µM GTP conditions, we next evaluated selected aminopyrimidines from our previous study (Fig. 1D). Although IC_50_ shifts between the two conditions are compound-specific, the overall SAR trends remained consistent. For instance, the IC_50_ of the lead compound **17** shifted from 86.2 nM in 20 µM guanosine to 163 nM in 200 µM GTP (Supplementary Table S1). This ∼2-fold decrease in apparent potency is attributable to the increased competition from the 10-fold higher concentration of the GTP cofactor relative to that of guanosine. In contrast, the IC_50_ of compound **11** was essentially unchanged between conditions, 207 nM versus 209 nM. These compound-specific responses indicate that cofactor conditions can influence compound–cofactor competition differently for each inhibitor, underscoring the need for physiologically calibrated assay conditions. Despite these compound-specific shifts, the rank-order of potency remained unchanged, with compound **17** emerging as the most potent inhibitor followed by compounds **11**, **12**, and **14**. Together, these data establish a physiologically calibrated assay that supports biologically meaningful SAR that can be used for design of downstream in-cell studies.

### Molecular beacon assay for high-throughput compound evaluation

To establish a high-throughput assay platform under the new conditions, we implemented a molecular beacon assay to evaluate a larger compound set.^23,35^ We first monitored the difference in dose-dependent responses between a radioanalytic assay and the molecular beacon assay (Supplementary Fig. S2). Overall SAR trends were preserved despite quantitative differences in potency. For example, compound **17** showed an IC_50_ of 163 nM in the radioanalytic assay and 472 nM in the molecular beacon assay, as observed previously.^23^ Compound **11** retained the same relative ranking, remaining less potent than compound **17** in both assay formats. Across the compound set, IC_50_ values from the molecular beacon assay were generally 4–10-fold higher than those obtained under the radioanalytic conditions (Supplementary Table S1). These shifts likely reflect the higher RNA concentration and relatively lower detection sensitivity of splicing events in the molecular beacon assay. Nevertheless, the optimized molecular beacon assay provided a practical high-throughput platform for screening of larger inhibitor libraries, and the most potent compounds were subsequently validated by gel-based radioanalytic assays.

### Structure-activity relationship (SAR) analysis of aminopyridine splicing inhibitors

To elucidate how each chemical moiety contributes to inhibition of C.a.mtLSU intron splicing, we performed an expanded SAR analysis of aminopyrimidines and aminopyridine derivatives (Fig. 3 and Supplementary Table S1). These analogues were rationally designed from the cryo-EM structure of the C.a.mtLSU intron bound to compound **11**, which defined the interactions required for aminopyrimidine binding. By distinguishing the functional groups that interact with the RNA from those pointing into solvent-accessible space, the structure suggested moieties that should be retained and others that were available for modification or expansion (Fig. 1B and Fig. 2). We first examined cyclic amine substitutions at the 4-position of the aminopyrimidine scaffold to probe the geometric requirements of the diamine moiety (Fig. 3A). Our previous work showed that linear amines are more potent than cyclic amines. However, in conventional medicinal chemistry, introducing conformational constraints and reducing entropic penalties by cyclizing amines often improves inhibitor potency.^36,37^ Because the previous study tested only a limited set of cyclic amines, it remained unclear whether cyclic amines are intrinsically suboptimal or only certain terminal amine geometries are suboptimal. We therefore synthesized 23 cyclic amine derivatives, compounds **1**–**10** and **25**–**37**, for systematic analysis.

**Figure 2.**
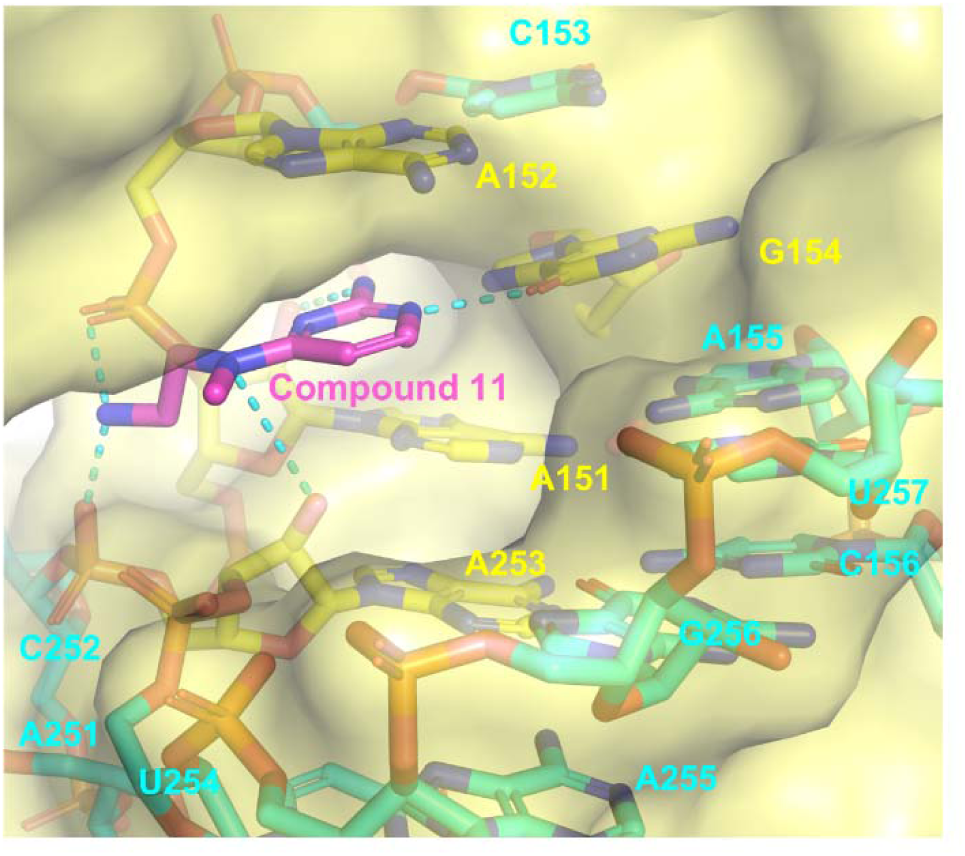
Compound 11 interaction with C.a.mtLSU intron nucleotides. Pink (compound **11**), yellow (nucleotides interacting with compound **11**), cyan (surrounding nucleotides which are not interacting with compound **11**)

**Figure 3.**
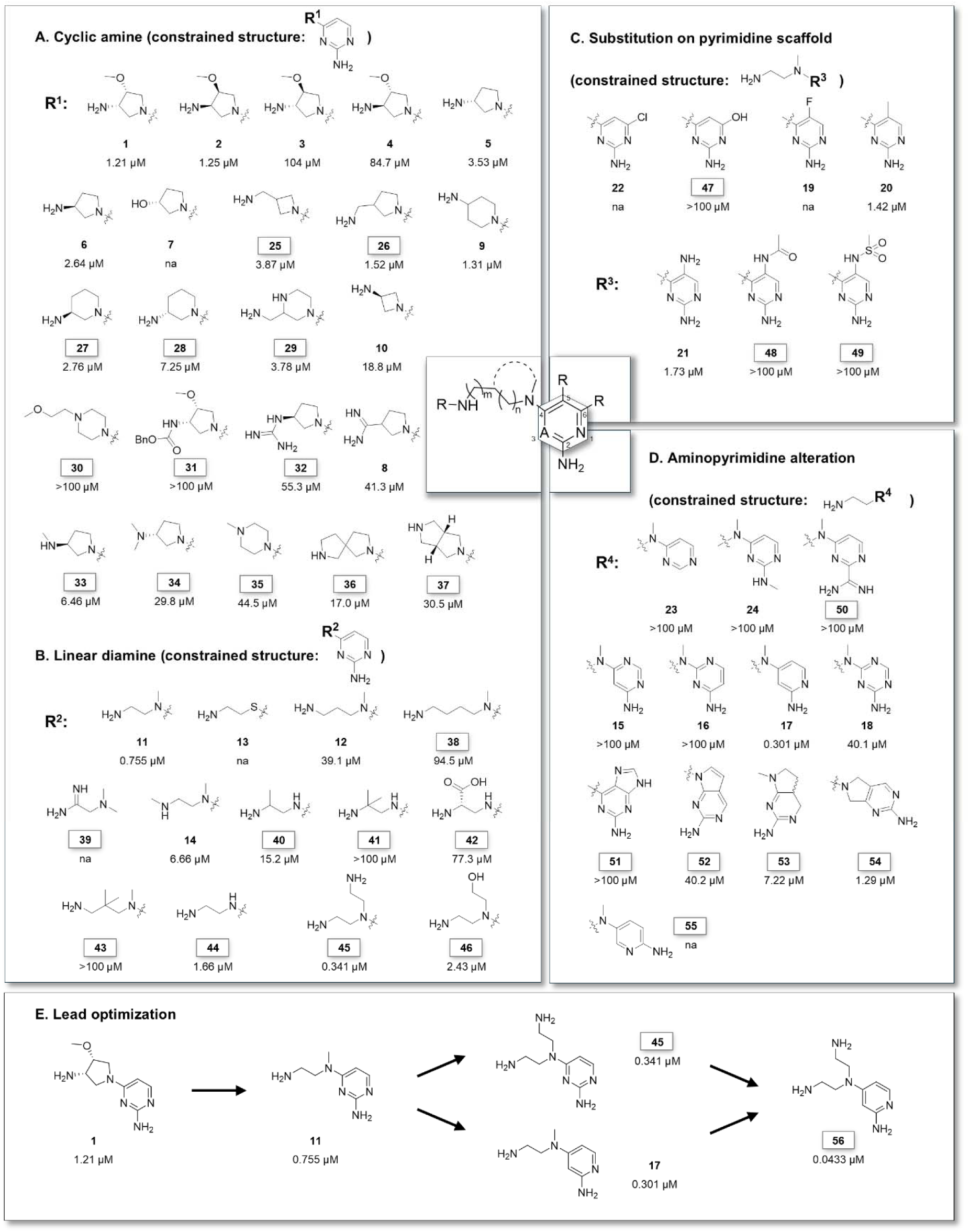
Aminopyrimidine and aminopyridine derivatives examined in the SAR study and their IC_50_ values in the molecular beacon assay. (A-D) Each moiety optimization based on SAR study. (E) Lead optimization discovered new lead compound **56** with IC_50_ of 43.3 nM. All compounds ID with box represents newly synthesized compound in this study.

Within this series of cyclic amine derivatives, the pyrimidine moiety was retained as 1,3-pyrimidin-2-amine. Additionally, all derivatives in this series retained their terminal amine which interacts with the phosphate backbone near A152 and A253 in the cryo-EM structure (Fig. 1B and Fig. 2). Compound **7** was an exception that has a terminal hydroxy group instead of amine as an inactive control. Compounds bearing a terminal primary amine consistently showed stronger inhibition than derivatives containing secondary or tertiary amines (Fig. 3A). Interestingly, low potency was observed for terminal primary amine derivatives that are trans conformers of compounds **3** and **4,** as observed in the previous study. Most terminal primary amine derivatives – including compounds **1**, **2**, **5**, **6**, **9**, and **25**–**29** – showed a comparable IC_50_ range except for compound **10,** despite their ring sizes and primary amine placement. A bulky substitution of the terminal amine (compounds **30** and **31**) or amidine substitution of the terminal primary amine (compounds **8** and **32**) resulted in even worse potency. When the terminal amine was embedded in a ring, IC_50_ increased, as in compounds **30** and **35**–**37**. However, no cyclic amine derivatives exceeded the potency of the linear diamine compound **11** (Fig. 3A and 3B). These data indicate that a terminal amine is indeed required for strong inhibition, but a highly constrained context is not needed. This behavior is consistent with an interaction between the terminal amines and the surrounding phosphate backbones, rather than a specific nucleobase. Indeed, the cyclic amine series explored here suggests that linear amines provide greater conformational plasticity, and may therefore be more compatible with the available RNA-ligand interface.

To expand our basic fragment-like scaffold into a more drug-like molecule, we explored growth vectors suggested by the cryo-EM structure, including the α and β carbons of the diamine linker, N4 position, and the aromatic C5 and C6 positions. These positions are solvent-exposed yet located near RNA residues that are capable of additional interactions, making them candidate sites for expansion (Fig. 2). We next performed SAR study within the linear diamine series while maintaining the pyrimidine moiety as 1,3-pyrimidin-2-amine (Fig. 3B). Consistent with our previous work, the aryl amine at the 4-position of the pyrimidine ring is required for activity, as thiol substitution in compound **13** completely abolished inhibition. Extension of the diamine linker to propylene or butylene, as in compounds **12** and **38**, substantially reduced potency.

Replacement of the terminal primary amine with amidine or secondary amine, as in compounds **14** and **39**, also abolished or reduced activity. We also tested whether additional substituents could be introduced into the ethylene or propylene linker based on apparent space in the binding pocket observed by cryo-EM (Fig. 1B and 2). Substitution at either the α- or β-position of the terminal amine was not tolerated, as observed for compounds **40**–**43**. Removal of the methyl group from the aryl amine in compound **44** caused only a modest decrease in activity. Notably, introduction of an additional *N*-ethyl amine in compound **45** increased potency whereas *N*-ethyl alcohol showed a 3-fold decrease in potency. Collectively, a *N*^4^,*N*^4^-bis(2-aminoethyl) group appears to be the preferred diamine moiety in the 4-position of the aminopyrimidine scaffold.

We next evaluated substitutions on the pyrimidine ring to evaluate the tolerance of ring modifications (Fig. 3C and 3D). We retained the 4-position of the aminopyrimidine moiety as a *N*^4^-(2-aminoethyl)-*N*^4^-methyl group. Only non-bulky electron-donating substituents at the 5-position, such as compounds **20** and **21**, were tolerated, although with modest reductions in potency (Fig. 3C). We then explored scaffold variation beyond the aminopyrimidine core (Fig. 3D). Amidine substitution of the aryl amine again caused a substantial loss of potency. Previous work had identified the 1-pyridine analog compound **17** as more potent than related 1,3-pyrimidine compounds. To explore this, we replaced the pyrimidine ring with a broader set of heteroaromatic purine-like scaffolds. All the tested scaffolds did not yield improved inhibitors although compound **54** showed a modest decrease in potency. Based on this analysis, pyridin-2-amine is the best scaffold in the heteroaromatic ring position.

The combined SAR study showed that the highest potency is achieved with a *N*^4^,*N*^4^-bis(2-aminoethyl) group at 4-position of pyrimidine and 1-pyridin-2-amine as the heterocycle (Fig. 3E). This result yielded a new lead compound **56** with IC_50_ of 43.3 nM. It is notable that compound **56** was substantially more potent than compounds **17** and **45**. This might be due to compatible placement of pyridine ring together with flexible bis-ethylamine strongly interacting with surrounding residues.

To validate these results using an orthogonal assay, selected compounds including the lead compounds **17** and **56** were further evaluated using a radioanalytic splicing assay conducted under the same GTP-based splicing conditions (Table 1). Overall SAR trends observed using both methods were the same, although IC_50_ values measured by the radioanalytic assay were generally 3–6-fold lower than those obtained with the molecular beacon assay. As mentioned above, higher IC_50_ values in the molecular beacon assay are likely attributable to lower sensitivity and higher RNA concentrations relative to those employed in the gel-based radioanalytic assay. Molecular beacon assays are also limited by the need for high concentrations of RNA, in which the lowest tested concentration of compounds is not in excess. Under the radioanalytic assay conditions, compound **17** exhibited an IC_50_ of 163 nM, whereas compound **56** showed an IC_50_ of 21.6 nM.

**Table 1.** IC_50_ of aminopyrimidines and aminopyridines in the molecular beacon and radioanalytic assays.

| Index | Compound | IC50 (µM)<br>at 200 µM GTP<sup>a</sup> | IC50 (µM)<br>at 200 µM GTP<sup>b</sup> |
|---|---|---|---|
| 1 | **1** | 1.21 | 0.270 |
| 2 | **2** | 1.25 | 0.290 |
| 3 | **7** | na | >100 |
| 4 | **9** | 1.31 | 0.420 |
| 5 | **11** | 0.755 | 0.209 |
| 6 | **12** | 39.1 | 9.73 |
| 7 | **14** | 6.66 | 1.96 |
| 8 | **17** | 0.472 | 0.163 |
| 9 | **21** | 1.73 | 0.519 |
| 10 | **26** | 1.52 | 0.533 |
| 11 | **44** | 1.66 | 0.438 |
| 12 | **45** | 0.341 | 0.178 |
| 13 | **46** | 2.43 | 0.706 |
| 14 | **53** | 7.22 | 2.13 |
| 15 | **56** | 0.0433 | 0.0216 |

<sup>a</sup> molecular beacon assay, <sup>b</sup> radioanalytic assay, na: not active

### Synergistic antifungal activity of aminopyridines with amphotericin B

To establish whether the mtLSU intron is the specific molecular target in cells, we first examined whether this intron actively splices in *Candida albicans* (strain SC5314) using RT-PCR (Fig. S3). Amplicons spanning the 5’exon-3’exon junction (lane 4) yield a cDNA product of ∼200 nt, which is the size expected for the ligated exons after splicing. These establish that the target intron is selectively excised and that the C.a.mtLSU intron splicing is active in fungal cells. We then evaluated whether this splicing event is affected upon administration of our aminopyridine compounds. *Candida albicans* was treated with lead inhibitors **17** and **56** and growth was monitored according to standardized CLSI protocols.^38^ Under standard growth conditions in RPMI, the compounds caused no detectable growth inhibition. This result is likely to reflect the poor uptake of aminopyridines due to electrostatic repulsion between the polycationic inhibitors and the negatively charged fungal membrane (data not shown). However, chemical permeabilization with benzalkonium chloride (BAC) resulted in a modest but mechanistically significant restoration of activity for the lead compound **56** (Supplementary Fig. S4). Even though this observed effect was marginal, it suggested that the initial lack of cellular efficacy could be due to delivery limitations rather than target irrelevance. This result prompted us to investigate whether co-administration with more potent clinical membrane-disrupting antifungal drugs might facilitate the aminopyridine uptake.

If successful, group I intron inhibition would represent a mechanistically distinct antifungal strategy relative to available FDA-approved antifungal agents, which act on the fungal membrane or cell wall by targeting ergosterol or β-1,3-glucan synthesis. Moreover, group I intron inhibition is expected to impair mitochondrial function, which is expected to reduce ATP-dependent drug efflux and thereby enhance the efficacy of conventional antifungal agents in a synergistic manner. To test whether our compounds might be efficacious in a synergistic fashion, we selected amphotericin B (AmpB) for combination studies because it is a broad-spectrum antifungal drug that disrupts fungal cell membrane integrity and could potentially alleviate the uptake limitation of these cationic compounds.

When compound **56** was combined with sublethal concentrations of amphotericin B (0.1–0.5 µg/mL), a pronounced growth inhibition of *C. albicans* was observed (Fig. 4A). Synergy was quantified using the Fractional Inhibitory Concentration (FIC) index, a metric that compares the potency of a drug combination to the individual activities of its components. The resulting FIC index of 0.33 falls below the established synergistic threshold of 0.5, indicating synergy between the aminopyridine **56** and the membrane-disrupting AmpB. This robust synergistic relationship demonstrates group I intron targeting represents an approach that is mechanistically orthogonal to traditional membrane-disrupting strategies. Furthermore, by impairing the splicing of essential mitochondrial genes, intron inhibition will disrupt mitochondrial metabolism, potentially reducing the ATP-dependent efflux.

**Figure 4.**
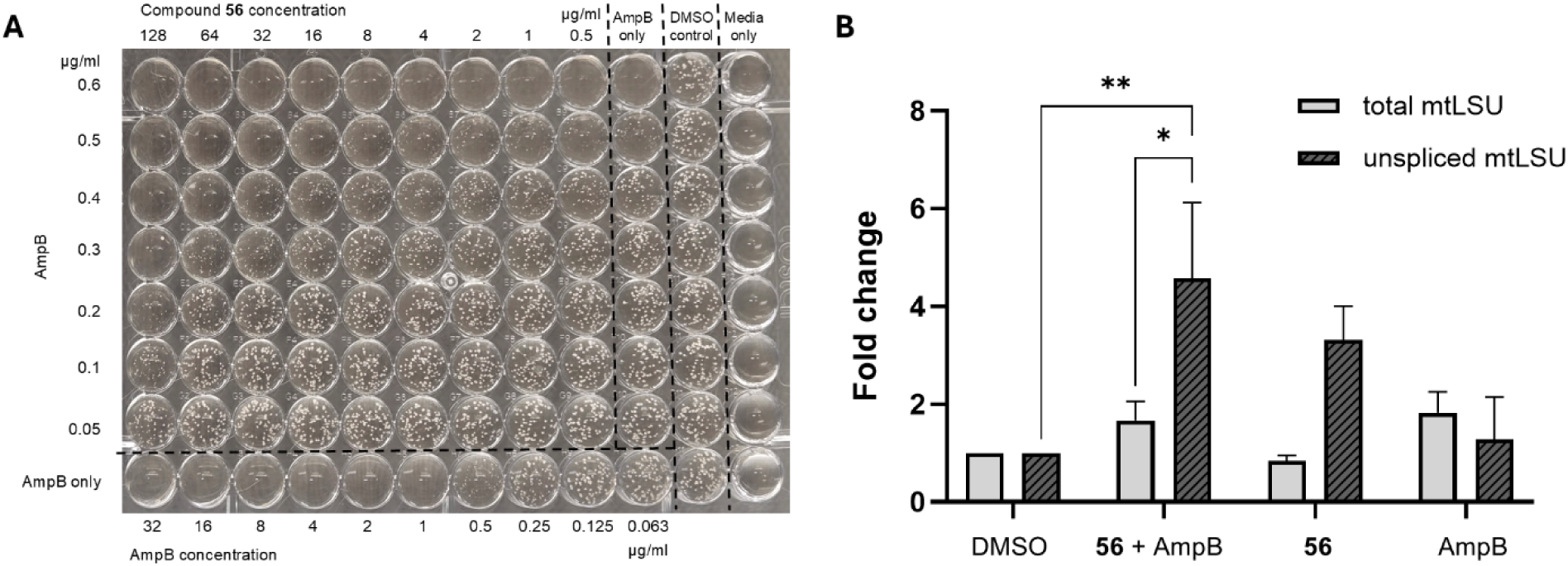
Synergistic antifungal activity of aminopyridine 56 with amphotericin B. (A) Antifungal assay of *C. albicans* SC5314 treated with aminopyridine **56** in combination with sublethal concentrations of amphotericin B (AmpB). The fractional inhibitory concentration (FIC) index for the combination is 0.33. (B) RT-qPCR analysis of C.a.mtLSU intron splicing under the indicated treatment conditions. The concentrations of **56** and AmpB were 64 μg/mL and 0.3 μg/mL, respectively. The relative fold change was normalized to housekeeping gene and DMSO.

To confirm target engagement and that the synergistic antifungal phenotype resulted from selective inhibition of the C.a.mtLSU intron splicing in cells, we quantified total and unspliced mtLSU transcript levels by RT-qPCR (Fig. 4B). We designed two primer sets to detect the enrichment of respective mtLSU. One amplifies the 5′ exon to report total mtLSU transcripts, while the other amplifies the 5′ exon–intron junction to report the unspliced precursor. While sublethal concentrations of AmpB alone had minimal impact on RNA processing, co-administration of AmpB and **56** caused a ∼4-fold accumulation of the unspliced mtLSU precursor relative to the DMSO control. Collectively, these data demonstrate that compound **56** disrupts the catalytic maturation of the mtLSU upon co-treatment with AmpB, validating the group I intron as the intracellular target of synergistic administration.

To our knowledge, this study provides not only the first functional evidence that pharmacological targeting of a fungal group I intron elicits an antifungal response in cells but also demonstrates that targeting fungal group I introns can produce synergistic antifungal effects with conventional antifungal drugs. These results establish group I intron as a clinically viable and mechanistically orthogonal target for antifungal drug development.

## Discussion

In this study, we establish that group I introns represent promising antifungal drug targets and that they also provide valuable platforms for developing an overall understanding of RNA-targeted small molecule discovery. Efforts to develop RNA targeting as a viable pharmacological strategy necessitate a greater diversity of validated targets, particularly those in which ligand binding is tightly coupled to a measurable biological function, together with SAR datasets that are large enough to yield generalizable design rules. Cases in which high resolution structures are available are even more valuable for establishing the first principles of RNA-small molecule recognition.

RNA molecules can sample multiple conformations and therefore establishing the pharmacologically relevant fold of an RNA is essential in order to conduct meaningful structure-function studies. This can involve optimizing parameters such as Mg² concentration, buffer conditions and temperature in the denaturation and refolding steps, so that screening assays reflect a biologically relevant conformation and competitive environment that is present in cells. In this work, we found that optimization of folding and enzymatic reaction conditions extended our ability to sensitively evaluate compound variations and ultimately obtain much more potent variants.

While many aspects of RNA and protein-targeted medicinal chemistry design are the same, this study suggests that “conventional wisdom” about structure-based design and optimization of inhibitor efficacy does not always apply to RNA. For example, rigidification via cyclization did not improve potency of our ligands, as a flexible linear diamine was more potent than every cyclic analogue. RNA forms stable, well-defined folds, but those folds can reorganize into new stable structures in response to ligand binding or to changes in the ionic or pH environment. This adaptive behavior, not a lack of structure, is the dynamic character of RNA, and it can favor a flexible ligand over a rigid one in engaging conformational flexibility in specific regions of the phosphate backbone. More generally, chemical features that are general rules in protein-directed design can be different for RNA as R-BIND suggests, so RNA-targeting compounds need design rules of their own, even though general principles such as stronger base/acid for tighter interaction still apply similarly. Guided by these insights, an aminopyridine bearing an *N□*,*N□*-bis(2-aminoethyl) group emerged as the most potent inhibitor, compound **56**, with an IC*□□*of 21.6 nM. It will be interesting to determine whether conformational plasticity of a ligand that also contains specific RNA interaction substituents will be a recurring requirement for engaging dynamic pockets within an RNA.

The main obstacle to turning *in vitro* potency into cellular activity is often delivery in RNA-targeting ligands due to the frequent presence of positively charged moieties. Despite systematic attempts to overcome this barrier, including spheroplast formation, non-ionic detergents, and prodrug masking (data not shown), only BAC-mediated permeabilization partially restored the cellular efficacy. These findings suggest that intracellular access, rather than target engagement, will be the principal barrier to many RNA-targeted compounds.

The fungal mitochondrion remains a largely unexplored pharmacological compartment, with no dedicated delivery study in *C. albicans* and no established route for importing antisense oligonucleotides across its membranes yet. Therefore, small molecules remain a particularly tractable modality for reaching mitochondrial RNA. The fungal mitochondrion is attractive both as a phylogenetically distinct target and as a pathway for disrupting resistance machinery of the cell. Strategies that exploit mitochondrial import, such as lipophilic cation conjugates and transporter-engaged prodrugs, can convert this biochemical potency into standalone cellular activity.

This study establishes RNA as a target that is not only orthogonal to protein targets but synergistic with protein or non-RNA-directed therapy, pointing to a strategy in which the two are engaged together to deepen efficacy or to counter resistance. The group I intron exemplifies this opportunity. The therapeutic value of combining orthogonal RNA- and protein-directed interventions is increasingly recognized across disease areas. For example, both chronic hepatitis B and castration-resistant prostate cancer are being pursued through complementary RNA- and protein-targeting approaches because protein-directed therapies alone have proven insufficient, although in those cases, the RNA component currently relies on antisense oligonucleotide targeting. Engaging RNA with a small molecule rather than a nucleic acid is particularly advantageous where the target is hard to reach, such as the central nervous system, where the blood-brain barrier confines oligonucleotides to intrathecal delivery, or disseminated viral infection, where broad biodistribution is required, since small molecules afford barrier penetration, broad tissue exposure, and oral dosing that nucleic-acid modalities largely cannot.

Taken together, this work provides one of the most extensive structurally rationalized SAR campaigns reported for an RNA target. This study establishes the group I intron as both a model system for RNA-targeted discovery and a clinically relevant, mechanistically distinct antifungal target. The molecular rules and the physiologically calibrated platform described here provide a foundation for RNA-targeted drug discovery. More broadly, we anticipate that combining calibrated functional assays and extensive SAR will accelerate the rational design of small molecules for other structured RNA targets.

## Methods

### *In vitro* transcription of C.a.mtLSU

All concentrations described below are final concentrations. The plasmid construct (pJH102, supplementary information) was obtained by point-mutation of previous reported plasmid (pLTS204) and linearized by using BamHI (NEB) according to manufacturer protocol. The C.a.mtLSU was then transcribed from the linearized plasmid using conditions adapted from those previously described.

1. In brief, for ^32^P-body labeled RNA, T7 RNA polymerase (P266L mutant) was added to mixture of 4 mM ATP, 4 mM CTP, 2 mM GTP, 0.8 mM UTP, and 50 µCi of [a-^32^P]-UTP (Revvity) in transcription buffer (40 mM Tris-Cl pH 8.0, 10 mM NaCl, 15 mM MgCl_2_, 2 mM spermidine, 0.01 % Triton X-100, and 10 mM DTT) and incubated at 37 °C for 1.5 h.
2. For unlabeled RNA, T7 RNA polymerase (P266L mutant) was added to mixture of 5 mM ATP, 5 mM CTP, 4 mM GTP, 5 mM UTP in transcription buffer (40 mM Tris-Cl pH 8.0, 10 mM NaCl, 20 mM MgCl_2_, 2 mM spermidine, 0.01 % Triton X-100, and 10 mM DTT) and incubated at 37 °C for 5 h.

The reaction was quenched with 2× urea loading buffer (10 M urea, 10 % sucrose, 0.2 % bromophenol blue, 0.2 % xylene cyanol, 40 mM Tris-Cl and 1 mM EDTA) and further purified on a 5% polyacrylamide (29:1) gel containing 8M urea. The corresponding precursor band was cut, crushed, and eluted in buffer (10 mM Na-MOPS pH 6.0, 100 mM NaCl, and 3 mM EDTA) at 4 °C for 3 h. The RNA transcript was collected by EtOH precipitation and stored in 6 mM Na-MES pH 6.0 at −20 °C.

## SPLICING ASSAY

### Radioanalytical splicing assay to monitor splicing

From the RNA stock solution prepared as described above, a ^32^P-body labeled C.a.mtLSU intron precursor construct at appropriate concentrations (see below) was prepared in 50 mM K-HEPES pH 7.5 and H_2_O, followed by incubation at 80 °C for 3 min to denature the RNA. The solution was then cooled to 25 °C for 30 min (2 °C/min). Then, 150 mM KCl and 30 mM MgCl_2_ were added to the solution, which was allowed to incubate for 5 min at 25 °C before the addition of DMSO or inhibitor compound in one of the two assays described below.

- Splicing kinetics: We conducted splicing time courses and measured *k*_obs_ values of different splicing conditions. DMSO was added to the folded intron precursor solution and incubated at 25 °C for 5 min with 5% final concentration of DMSO. As soon as guanosine or GTP added to RNA (10 nM), timepoints of 0, 1.5, 3, 5, 7.5, 10, 20, 30, 45, 60 min were taken.
- Quantitative IC_50_ determination: Compounds were initially diluted to a concentration of 6 mM, 2 mM, 600 µM, 200 µM, 60 µM, 20 µM, 6 µM, 2 µM, 600 nM, 200 nM, 60 nM, 20 nM in DMSO prior to addition to the RNA (2 nM). A corresponding amount of compound (300 µM, 100 µM, 30 µM, 10 µM, 3 µM, 1 µM, 300 nM, 100 nM, 30 nM, 10nM, 3 nM, 1 nM, final concentration) was added to the folded intron precursor solution and incubated at 25 °C for 10 min with 5% final concentration of DMSO. In the final step, 200 µM GTP was added to initiate the splicing reaction. The splicing reaction was incubated at 25 °C for 10 minutes and then quenched with ice-cold 2× formamide loading buffer (72 % (v/v) formamide, 10 % sucrose, 0.2 % bromophenol blue, and 50 mM EDTA). Samples were then loaded onto 5% polyacrylamide (29:1) gel containing 8 M urea to resolve individual bands.

### Data analysis of radioanalytic assay

Gels were dried at 80 °C for 30 min and exposed to a phosphor screen overnight. The screen was then scanned using a Typhoon RGB Biomolecular imager (Cytiva). Individual bands were quantified by ImageQuant 8.2 software (GE Healthcare). Band intensity was corrected using the uridine content of each corresponding RNA fragment, followed by normalization by total number of RNA (precursor, intermediate, and linear intron). All assays were performed in triplicate.

- Data were plotted with an equation below to obtain a *k*_obs_ value.

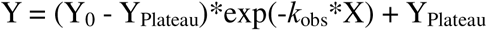

with following definitions; Y = Precursor (%), X = Time (min)

- The half-maximal inhibitory concentration (IC_50_) was determined by fitting the following equation:

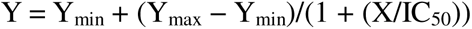

with following definitions; Y = Linear intron (%), X = Concentration (µM)

### Molecular beacon assay to monitor splicing inhibition

Compound activity was assessed using a molecular beacon-based fluorescence assay in a 384-well format (40 µL final reaction volume). The molecular beacon was designed as previously described, except that the fluorophore–quencher pair was changed to Texas Red and Black Hole Quencher 2 (BHQ2) to minimize interference from the autofluorescence of small-molecule compounds (MB-TxR-JH102-1: 5′-TxR-CCAGGTACCCTGTTATCCCTAGCCCTGG-BHQ2-3′).

For IC_50_ determination, compounds were prepared as a serial dilution in DMSO and dispensed into wells with H O to give final concentrations ranging from 1 nM to 300 µM (1, 3, 10, 30, 100, 300 nM and 1, 3, 10, 30, 100, 300 µM) at a constant final DMSO concentration of 5 %. RNA (20 nM final) was folded by mixing with 50 mM HEPES buffer (pH 7.5), denaturing at 80 °C for 5 min, and slow cooling at room temperature for 30 min; 150 mM KCl and 3 mM MgCl_2_ were then added, followed by incubation at room temperature for 5 min. The folded RNA was added to the compound-containing wells and incubated at 25 °C for 10 min. Splicing was initiated by addition of 200 µM GTP, and the reaction was incubated at 25 °C for 10 min before quenching with 10 mM EDTA. The molecular beacon (5 nM) was then added, and plates were incubated at 42 °C for 30 min followed by room temperature for 30 min. Reactions lacking compound (DMSO vehicle) or lacking RNA (H O) served as positive and negative controls, respectively.

Fluorescence was measured on a BioTek Neo plate reader in monochromator mode (excitation 580/10 nm, emission 620/10 nm). Percent inhibition was calculated as below:

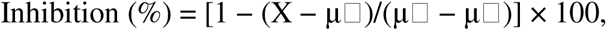

where X = sample fluorescence, μ□ = mean fluorescence of the positive controls, μ□ = mean fluorescence of the negative controls.

## FUNGAL GROWTH ASSAY

### Compound–drug co-administration growth assay

Fungal growth inhibition was determined by broth microdilution in RPMI-1640. Test compounds were serially diluted two-fold across nine points in DMSO from a 25.6 mg/mL stock, and the antifungal drug (AmpB) was prepared at fixed sub-inhibitory concentrations (120, 100, 80, 60, 40, 20, and 10 µg/mL). Compound and drug preparations were each diluted 1:100 respectively, into RPMI-1640 prior to addition to cells, and the final DMSO concentration in the assay was 1 %. *C. albicans* colonies were inoculated in medium, and the cell density was determined using an automated cell counter and adjusted to 1–5 × 10□ CFU/mL. The suspension was serially diluted in RPMI-1640, and 50 µL was dispensed per well and combined 1:1 with 50 µL of the compound/drug/medium dilutions in a 96-well plate to give a final inoculum of approximately 1–5 × 10³ CFU/mL. Wells containing medium alone served as no-growth control while wells with DMSO served as the negative control. Plates were incubated at 37 °C for 24 h, and growth was assessed visually to determine the minimum concentration at which growth was inhibited.

### Detergent-mediated permeabilization

To evaluate the effect of membrane permeabilization on compound activity, cells were pretreated with benzalkonium chloride (BAC) prior to the compounds treatment. Following inoculum preparation as described above, cells were pelleted (2,000 × g, 5 min), washed twice with PBS, and resuspended in PBS containing BAC at the indicated concentration. After incubation at room temperature for 1 h, cells were pelleted, washed twice with PBS, and resuspended in RPMI-1640. Cell density was then determined with an automated cell counter, and the suspension was diluted and used in the growth inhibition assay as described above.

### Cell culture and harvest to extract total RNA

A single colony was inoculated into RPMI-1640 (pH 7.0) medium and grown overnight at 30 °C with shaking at 250 rpm. The saturated overnight culture was diluted with RPMI-1640 to an OD_600_ of 0.2 in a final volume of 5 mL and grown to mid-log phase (OD_600_ ≈ 0.5) at 30 °C with shaking at 250 rpm.

1. Cell harvest for RT-PCR: Cells were harvested by centrifugation at 3,000 rpm for 5 min, and the supernatant was discarded. The pellet was resuspended in 1 mL of ice-cold H□O, transferred to a screw-cap tube, and collected by centrifugation at 5,000 × g for 5 min. After removal of the supernatant, the cell pellet was flash-frozen in liquid N□ and stored at −80 °C until RNA extraction.
2. Compound treatment: Mid-log cultures were aliquoted into 5 mL each and treated with compound dissolved in DMSO, keeping the final DMSO concentration at 1 % in RPMI-1640 (e.g., 64 µg/mL final from a 12.8 mg/mL stock). For co-administration, the test compound and antifungal drug were added together at their respective concentrations while holding total DMSO constant, and 1 % DMSO alone served as the negative control. Cultures were incubated at 30 °C with shaking at 250 rpm for 3 h and harvested by centrifugation at 2,500 × g for 5 min. Cell pellets were resuspended in 0.5 mL of buffer (1 M sorbitol, 20 mM potassium phosphate, 50 mM EDTA, 0.1 % BME) and spheroplasted with zymolyase (0.25 U per 10□ cells) at 30 °C for 20 min. Spheroplasts were gently collected by centrifugation at 1,000 × g for 5 min at room temperature to avoid lysis, flash-frozen in liquid N□, and stored at −80 °C until RNA extraction.

### RNA extraction

Total RNA was extracted using the E.Z.N.A. kit according to the manufacturer’s protocol. To remove residual DNA, each RNA sample (50 µL) was treated with 6 µL of RQ1 DNase buffer, 6 µL of RQ1 DNase, and 1 µL of RNase inhibitor, and incubated at 30 °C for 10 min. The RNA was then recovered by EtOH precipitation. The RNA pellet was air-dried for 10 min and resuspended in 30 µL of ME buffer supplemented with RNase inhibitor (1:100 v/v). RNA concentration was determined by NanoDrop.

### Reverse transcription (RT)

First-strand cDNA was synthesized from 200 ng of total RNA using SuperScript III reverse transcriptase (Invitrogen) with random hexamer primers, according to the manufacturer’s protocol. Following primer annealing (65 °C, 5 min), reverse transcription was carried out in a 20 µL reaction at 25 °C for 5 min, 50 °C for 1 h, and 70 °C for 15 min. A parallel no-RT control lacking reverse transcriptase was included for each sample. The RNA template was then degraded with RNase H, RNase A, and RNase T1 (0.5 µL each) at 37 °C for 30 min, and cDNA was purified using AMPure XP beads at a 1.8× ratio and eluted in 32 µL of nuclease-free H O.

### PCR and agarose gel electrophoresis

cDNA was amplified using Q5 High-Fidelity DNA Polymerase (NEB) in a 25 µL reaction containing 1× Q5 reaction buffer, 200 µM dNTPs, 0.5 µM each of the forward and reverse primers (supplementary information), 0.02 U/µL Q5 polymerase, and 1 µL of RT template. pJH102 plasmid (10 ng/µL) and nuclease-free H O served as positive and negative template controls, respectively. Thermocycling was performed with an initial denaturation at 98 °C for 20 s, followed by 35 cycles of 98 °C for 10 s, 60 °C for 30 s, and 72 °C for 20–40 s, a final extension at 72 °C for 20 s. Amplicons were resolved on a 1 % agarose gel prepared in 1× TAE buffer (40 mM Tris, 20 mM acetic acid, and 1 mM EDTA) containing GelRed. Bands were visualized by fluorescence imaging (Typhoon).

### Quantitative PCR (qPCR)

Transcript levels of compound-treated cells were quantified by qPCR using LightCycler® 480 SYBR Green I Master (Roche). Each 10 µL reaction contained 1× SYBR Green Master Mix, 1 µM each of the forward and reverse primers (Supplementary information), and cDNA template, assembled in a white 384-well plate. qPCR was performed on a real-time PCR instrument with an initial denaturation at 95 °C for 10 min, followed by 40 cycles of 95 °C for 10 s, 62 °C for 20 s, and 72 °C for 30 s. Melt-curve analysis was performed by denaturation at 95 °C for 10 s, re-annealing at 62 °C for 10 s, and a stepwise ramp from 62 °C to 95 °C (0.5 °C per 30 s) with fluorescence measured at each step.

Data were analyzed by the ΔΔC_q_ method. Reactions were retained only when negative controls (no-reverse transcriptase) gave C_q_ ≥ 35 and RT samples gave C_q_ between 10 and 30, and technical replicates were averaged. Transcript levels were normalized to the *ACT1* reference gene, and fold changes in test-compound-treated samples were calculated relative to the untreated control as below:

Fold change = 2^(−ΔΔCq)^,

where ΔΔC_q_ = (C_q,target gene_ – C_q,ACT1_)_compound_ − (C_q,target gene_ – C_q,ACT1_)_DMSO_. Total (unspliced + spliced) and unspliced mtLSU transcripts were quantified using distinct primer sets, and fold changes for total versus unspliced transcripts were compared by unpaired t-test.

## Acknowledgements

A.M.P. is an investigator of the HHMI. We are grateful to M. Luo and H. Jang for experimental discussion of RT-qPCR and for proofreading the manuscript.

## Author contributions

A.M.P. and J.H. designed the research. J.H. performed all experiments except for chemical synthesis. L.J.S. and T.S. conducted chemical synthesis. L.J.S. and M.C.V.Z. designed small molecules. A.M.P. and J.H. wrote the manuscript.

## References

1. Warner, K.D., Hajdin, C.E., and Weeks, K.M. (2018). Principles for targeting RNA with drug-like small molecules. Nat Rev Drug Discov 17, 547–558. 10.1038/nrd.2018.93.

2. Falese, J.P., Donlic, A., and Hargrove, A.E. (2021). Targeting RNA with small molecules: from fundamental principles towards the clinic. Chem Soc Rev 50, 2224–2243. 10.1039/d0cs01261k.

3. Velagapudi, S.P., Costales, M.G., Vummidi, B.R., Nakai, Y., Angelbello, A.J., Tran, T., Haniff, H.S., Matsumoto, Y., Wang, Z.F., Chatterjee, A.K., et al. (2018). Approved Anti-cancer Drugs Target Oncogenic Non-coding RNAs. Cell Chem Biol 25, 1086–1094.e1087. 10.1016/j.chembiol.2018.05.015.

4. Schwanhausser, B., Busse, D., Li, N., Dittmar, G., Schuchhardt, J., Wolf, J., Chen, W., and Selbach, M. (2011). Global quantification of mammalian gene expression control. Nature 473, 337–342. 10.1038/nature10098.

5. Hausser, J., Mayo, A., Keren, L., and Alon, U. (2019). Central dogma rates and the trade-off between precision and economy in gene expression. Nat Commun 10, 68. 10.1038/s41467-018-07391-8.

6. Garner, A.L. (2023). Contemporary Progress and Opportunities in RNA-Targeted Drug Discovery. ACS Medicinal Chemistry Letters 14, 251–259. 10.1021/acsmedchemlett.3c00020.

7. Butcher, S.E., and Pyle, A.M. (2011). The molecular interactions that stabilize RNA tertiary structure: RNA motifs, patterns, and networks. Acc Chem Res 44, 1302–1311. 10.1021/ar200098t.

8. Ganser, L.R., Kelly, M.L., Herschlag, D., and Al-Hashimi, H.M. (2019). The roles of structural dynamics in the cellular functions of RNAs. Nat Rev Mol Cell Biol 20, 474–489. 10.1038/s41580-019-0136-0.

9. Olenginski, L.T., Wierzba, A.J., Laursen, S.P., and Batey, R.T. (2026). Designing small molecules targeting a cryptic RNA binding site through base displacement. Nat Chem Biol 22, 1004–1013. 10.1038/s41589-025-02018-8.

10. Stelzer, A.C., Frank, A.T., Kratz, J.D., Swanson, M.D., Gonzalez-Hernandez, M.J., Lee, J., Andricioaei, I., Markovitz, D.M., and Al-Hashimi, H.M. (2011). Discovery of selective bioactive small molecules by targeting an RNA dynamic ensemble. Nat Chem Biol 7, 553–559. 10.1038/nchembio.596.

11. Velagapudi, S.P., Gallo, S.M., and Disney, M.D. (2014). Sequence-based design of bioactive small molecules that target precursor microRNAs. Nat Chem Biol 10, 291–297. 10.1038/nchembio.1452.

12. Howe, J.A., Wang, H., Fischmann, T.O., Balibar, C.J., Xiao, L., Galgoci, A.M., Malinverni, J.C., Mayhood, T., Villafania, A., Nahvi, A., et al. (2015). Selective small-molecule inhibition of an RNA structural element. Nature 526, 672–677. 10.1038/nature15542.

13. Donlic, A., Morgan, B.S., Xu, J.L., Liu, A., Roble, C., Jr., and Hargrove, A.E. (2018). Discovery of Small Molecule Ligands for MALAT1 by Tuning an RNA-Binding Scaffold. Angew Chem Int Ed Engl 57, 13242–13247. 10.1002/anie.201808823.

14. Fedorova, O., Jagdmann, G.E., Jr., Adams, R.L., Yuan, L., Van Zandt, M.C., and Pyle, A.M. (2018). Small molecules that target group II introns are potent antifungal agents. Nat Chem Biol 14, 1073–1078. 10.1038/s41589-018-0142-0.

15. Childs-Disney, J.L., Yang, X., Gibaut, Q.M.R., Tong, Y., Batey, R.T., and Disney, M.D. (2022). Targeting RNA structures with small molecules. Nat Rev Drug Discov 21, 736–762. 10.1038/s41573-022-00521-4.

16. Ratni, H., Ebeling, M., Baird, J., Bendels, S., Bylund, J., Chen, K.S., Denk, N., Feng, Z., Green, L., Guerard, M., et al. (2018). Discovery of Risdiplam, a Selective Survival of Motor Neuron-2 (SMN2) Gene Splicing Modifier for the Treatment of Spinal Muscular Atrophy (SMA). Journal of Medicinal Chemistry 61, 6501–6517. 10.1021/acs.jmedchem.8b00741.

17. Maucort, C., Vo, D.D., Aouad, S., Charrat, C., Azoulay, S., Di Giorgio, A., and Duca, M. (2021). Design and Implementation of Synthetic RNA Binders for the Inhibition of miR-21 Biogenesis. ACS Medicinal Chemistry Letters 12, 899–906. 10.1021/acsmedchemlett.0c00682.

18. Connelly, C.M., Numata, T., Boer, R.E., Moon, M.H., Sinniah, R.S., Barchi, J.J., Ferre-D’Amare, A.R., and Schneekloth, J.S., Jr. (2019). Synthetic ligands for PreQ1 riboswitches provide structural and mechanistic insights into targeting RNA tertiary structure. Nature Communications 10, 1501. 10.1038/s41467-019-09493-3.

19. Donlic, A., Swanson, E.G., Chiu, L.Y., Wicks, S.L., Juru, A.U., Cai, Z., Kassam, K., Laudeman, C., Sanaba, B.G., Sugarman, A., et al. (2022). R-BIND 2.0: An Updated Database of Bioactive RNA-Targeting Small Molecules and Associated RNA Secondary Structures. ACS Chem Biol 17, 1556–1566. 10.1021/acschembio.2c00224.

20. Morgan, B.S., Forte, J.E., Culver, R.N., Zhang, Y., and Hargrove, A.E. (2017). Discovery of Key Physicochemical, Structural, and Spatial Properties of RNA-Targeted Bioactive Ligands. Angewandte Chemie International Edition 56, 13498–13502. 10.1002/anie.201707641.

21. Duca, M. (2024). Small molecule approaches to targeting RNA. Nature Reviews Chemistry 8, 120–135. 10.1038/s41570-023-00569-9.

22. Liu, T., and Pyle, A.M. (2024). Highly Reactive Group I Introns Ubiquitous in Pathogenic Fungi. J Mol Biol 436, 168513. 10.1016/j.jmb.2024.168513.

23. Liu, T., Xu, L., Chung, K., Sisto, L.J., Hwang, J., Zhang, C., Van Zandt, M.C., and Pyle, A.M. (2025). Molecular insights into de novo small-molecule recognition by an intron RNA structure. Proc Natl Acad Sci U S A 122, e2502425122. 10.1073/pnas.2502425122.

24. Cech, T.R., Zaug, A.J., and Grabowski, P.J. (1981). In vitro splicing of the ribosomal RNA precursor of Tetrahymena: involvement of a guanosine nucleotide in the excision of the intervening sequence. Cell 27, 487–496. 10.1016/0092-8674(81)90390-1.

25. Adams, P.L., Stahley, M.R., Kosek, A.B., Wang, J., and Strobel, S.A. (2004). Crystal structure of a self-splicing group I intron with both exons. Nature 430, 45–50. 10.1038/nature02642.

26. Treiber, D.K., and Williamson, J.R. (1999). Exposing the kinetic traps in RNA folding. Curr Opin Struct Biol 9, 339–345. 10.1016/s0959-440x(99)80045-1.

27. Zhuang, X., Bartley, L.E., Babcock, H.P., Russell, R., Ha, T., Herschlag, D., and Chu, S. (2000). A single-molecule study of RNA catalysis and folding. Science 288, 2048–2051. 10.1126/science.288.5473.2048.

28. Haugen, P., Simon, D.M., and Bhattacharya, D. (2005). The natural history of group I introns. Trends Genet 21, 111–119. 10.1016/j.tig.2004.12.007.

29. Pang, K.C., Frith, M.C., and Mattick, J.S. (2006). Rapid evolution of noncoding RNAs: lack of conservation does not mean lack of function. Trends Genet 22, 1–5. 10.1016/j.tig.2005.10.003.

30. Hezroni, H., Koppstein, D., Schwartz, M.G., Avrutin, A., Bartel, D.P., and Ulitsky, I. (2015). Principles of long noncoding RNA evolution derived from direct comparison of transcriptomes in 17 species. Cell Rep 11, 1110–1122. 10.1016/j.celrep.2015.04.023.

31. Fisher, M.C., Alastruey-Izquierdo, A., Berman, J., Bicanic, T., Bignell, E.M., Bowyer, P., Bromley, M., Bruggemann, R., Garber, G., Cornely, O.A., et al. (2022). Tackling the emerging threat of antifungal resistance to human health. Nat Rev Microbiol 20, 557–571. 10.1038/s41579-022-00720-1.

32. Fisher, M.C., and Denning, D.W. (2023). The WHO fungal priority pathogens list as a game-changer. Nat Rev Microbiol 21, 211–212. 10.1038/s41579-023-00861-x.

33. Russell, R., Zhuang, X., Babcock, H.P., Millett, I.S., Doniach, S., Chu, S., and Herschlag, D. (2002). Exploring the folding landscape of a structured RNA. Proceedings of the National Academy of Sciences of the United States of America 99, 155–160. 10.1073/pnas.221593598.

34. Park, J.O., Rubin, S.A., Xu, Y.F., Amador-Noguez, D., Fan, J., Shlomi, T., and Rabinowitz, J.D. (2016). Metabolite concentrations, fluxes and free energies imply efficient enzyme usage. Nat Chem Biol 12, 482–489. 10.1038/nchembio.2077.

35. Omran, Q.Q., Fedorova, O., Liu, T., and Pyle, A.M. (2022). A molecular beacon assay for monitoring RNA splicing. Nucleic Acids Res 50, e74. 10.1093/nar/gkac242.

36. DeLorbe, J.E., Clements, J.H., Teresk, M.G., Benfield, A.P., Plake, H.R., Millspaugh, L.E., and Martin, S.F. (2009). Thermodynamic and structural effects of conformational constraints in protein-ligand interactions. Entropic paradoxy associated with ligand preorganization. J Am Chem Soc 131, 16758–16770. 10.1021/ja904698q.

37. Zhang, L., Zhang, B., Tian, L., Xia, J., Wang, M., Chang, H., Hong, T., Liang, C., and Zhao, M. (2025). Conformationally restricted pleuromutilins via a pyridinyl-amide tether bias peptidyl transferase center engagement enabling broad-spectrum potency against multidrug-resistant bacteria. Eur J Med Chem 302, 118316. 10.1016/j.ejmech.2025.118316.

38. Clinical and Laboratory Standards Institute (2017). Reference Method for Broth Dilution Antifungal Susceptibility Testing of Yeasts.

