## Supplemental information for "RNA-targeted aminopyridine compounds provide a new antifungal strategy"

**Contents**

**Figure S1**	**2**

**Figure S2**	**3**

**Figure S3**	**4**

**Figure S4**	**5**

**Table S1**	**6**

**Methods**	**16**

**Chemical Synthesis and Characterization**	**17**

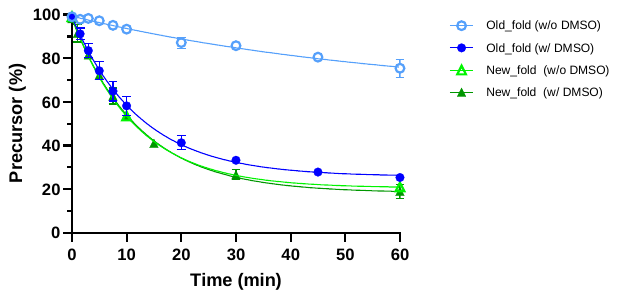

**Figure S1.** C.a.mtLSU splicing timecourse in different folding conditions. Previous folding: denaturing RNA at 90 °C for 1 min, folding RNA with cooling at 25 °C for 2 min. New folding: denaturing RNA at 80 °C for 3 min, folding RNA with cooling to 25 °C for 30 min (2 °C/min). Final DMSO concentration is 5 %.

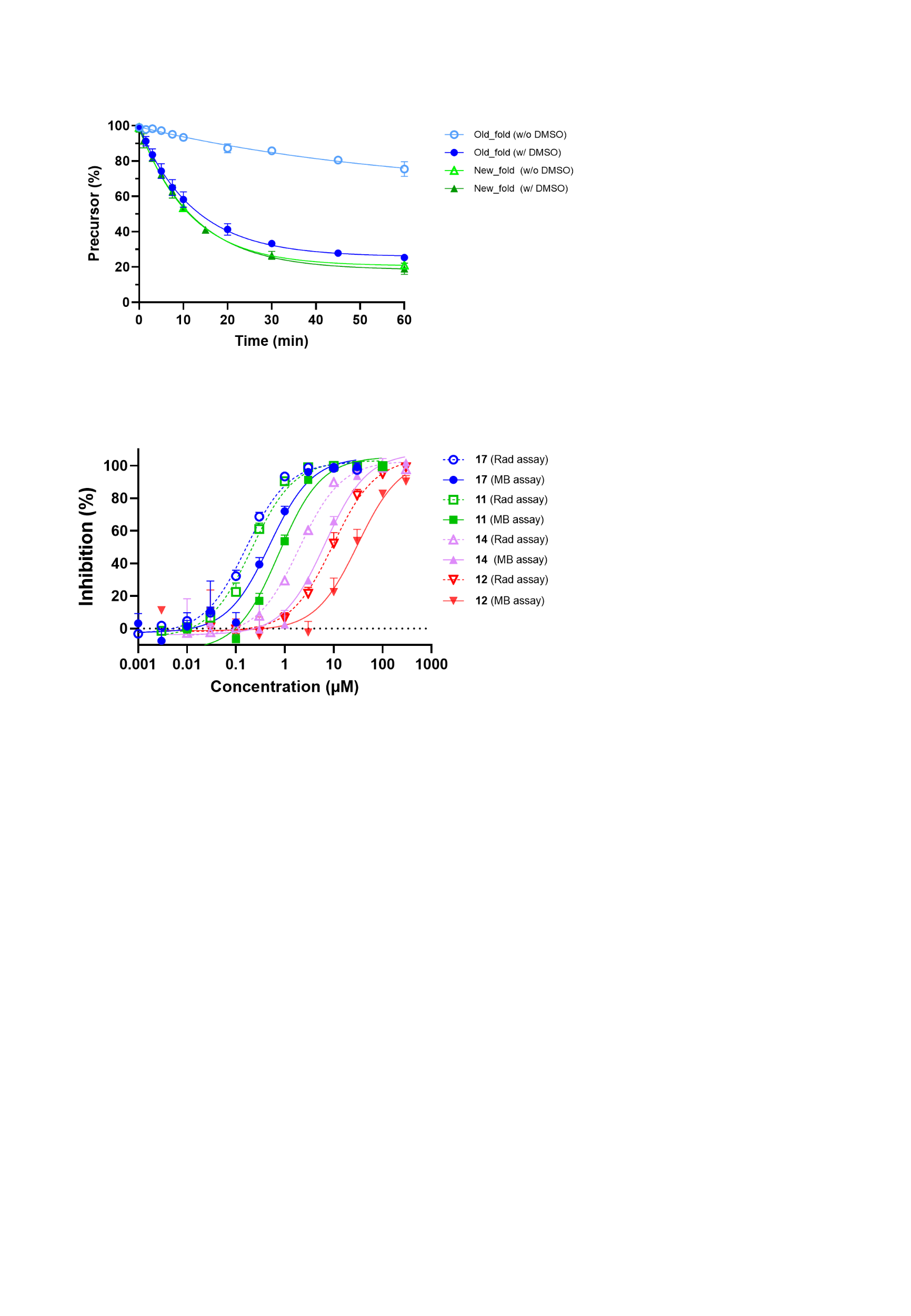

**Figure S2.** Dose–response curves for aminopyrimidine-mediated splicing inhibition under two different assays, radioanalytic assay (Rad) and molecular beacon assay (MB). Dotted lines represent radioanalytic assay and solid lines represent molecular beacon assay. IC50 of each compounds are 0.163 μM (Rad), 0.490 μM (MB) for **17**, 0.209 μM (Rad), 0.755 μM (MB) for **11**, 1.96 μM (Rad), 6.66 μM (MB) for **14**, 9.24 μM (Rad), 31.2 μM (MB) for **12**, respectively.

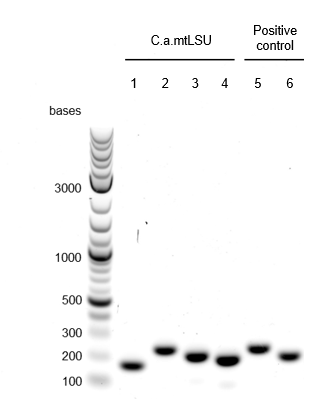

**Figure S3.** RT-PCR analysis of mitochondrial mtLSU transcripts from C. albicans SC5314. Lanes 1–4 probe the C.a. mtLSU; lanes 5–6 are housekeeping controls. Lane 1, 5′ exon (total mtLSU transcript); lane 2, unspliced mtLSU (spanning 5’ exon–intron); lane 3, ligated exon (spanning 5′ exon–3′ exon); lane 4, ligated exon (spanning 5′ exon–3′ exon junction); lane 5, ACT1; lane 6, PGK1. Detection of the ~200 nt ligated exon products (lane 3 and 4) confirms that mtLSU splicing occurs in cells

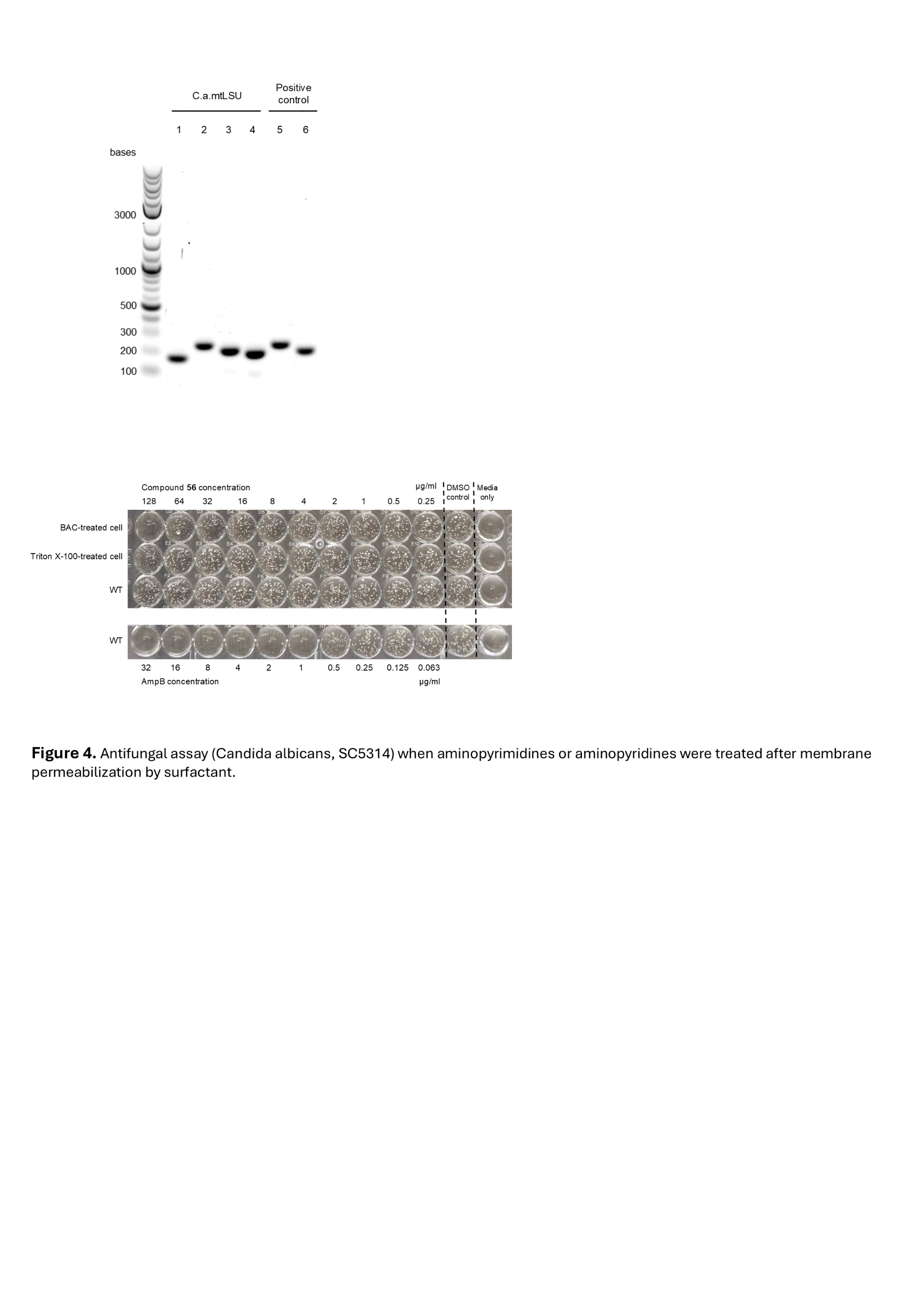

**Figure S4.** Fungal growth inhibition of aminopyrimidine and aminopyridine compounds following membrane permeabilization. Growth of *C. albicans* SC5314 was monitored after treatment with aminopyridine **56** under untreated conditions or following pretreatment with 1 mg/mL Triton X-100 or 20 μg/mL benzalkonium chloride (BAC).

**Table S1**. Dose-dependent inhibition of aminopyrimidines and aminopyridines in molecular beacon assay and radioanalytic assay

| **Index** | **Compound ID** | | **Chemical structure** | **IC50** | |
| --- | --- | --- | --- | --- | --- |
|  |  |  |  | **Molecular beacon assay** | **Radioanalytic assay** |
| 1 | **1** | **628** | 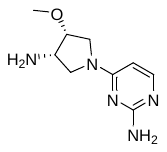 | 1.21 μM  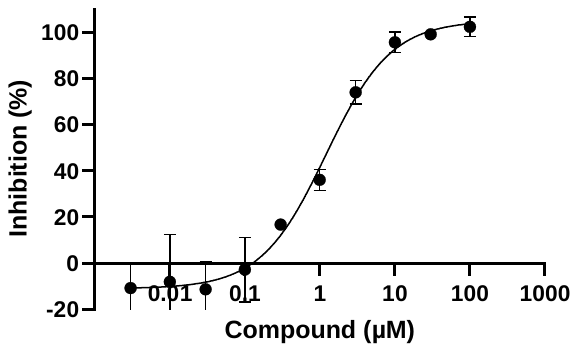 | 0.270 μM  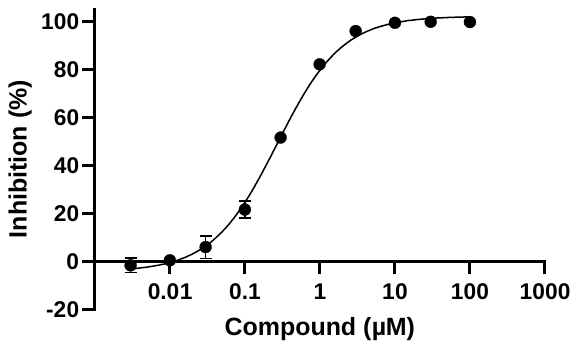 |
| 2 | **2** | **626** | 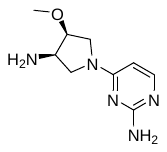 | 1.26 μM  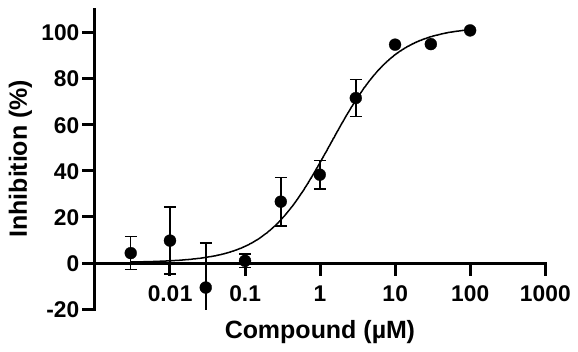 | 0.290 μM  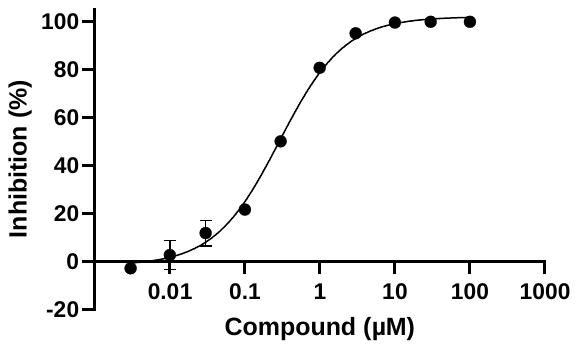 |
| 3 | **3** | **614** | 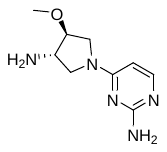 | 104 μM  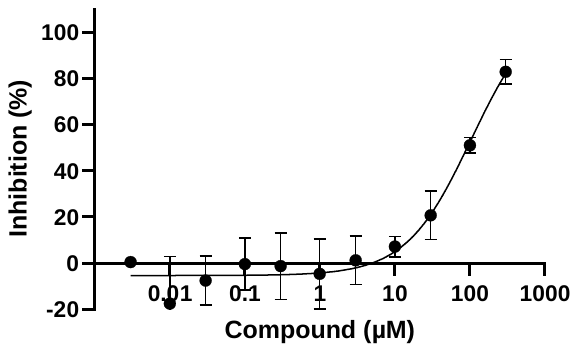 |  |
| 4 | **4** | **617** | 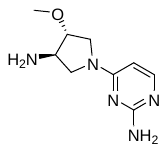 | 84.7 μM  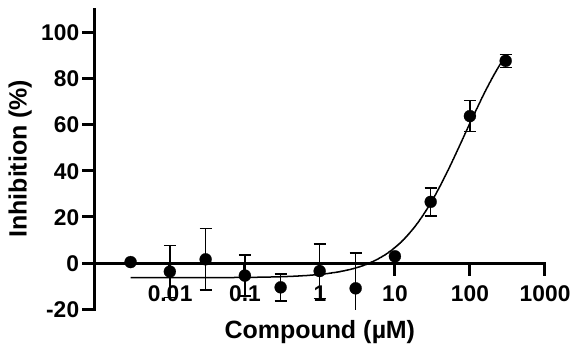 |  |
| 5 | **5** | **613** | 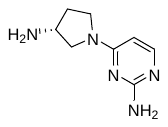 | 3.53 μM  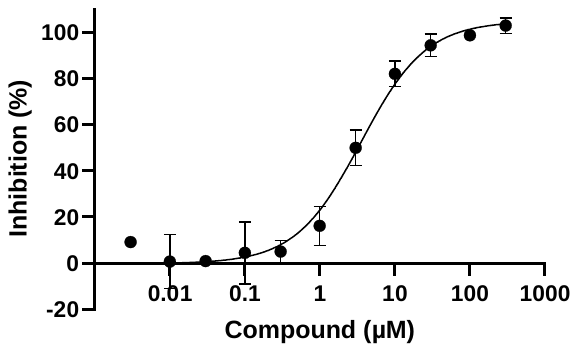 |  |
| 6 | **6** | **615** | 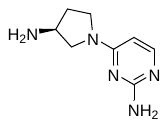 | 2.64 μM  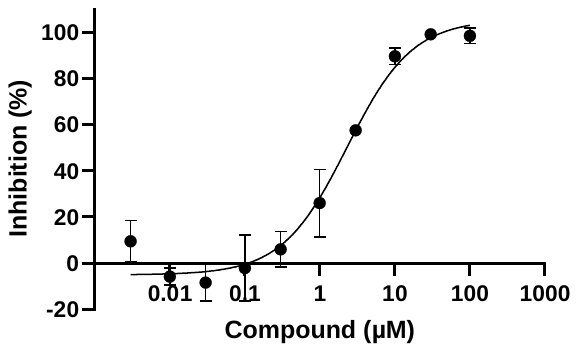 |  |
| 7 | **7** | **618** | 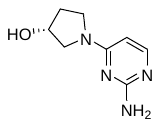 | Not active  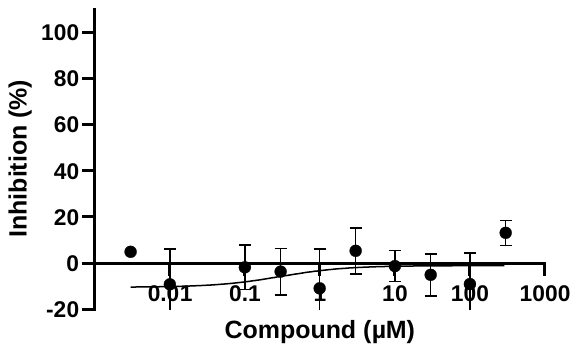 | >100 μM  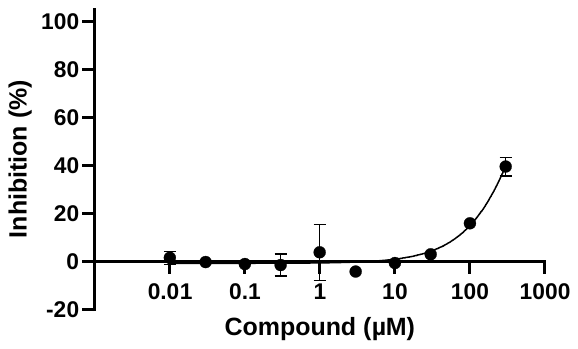 |
| 8 | **25** | **711** | 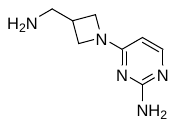 | 3.87 μM  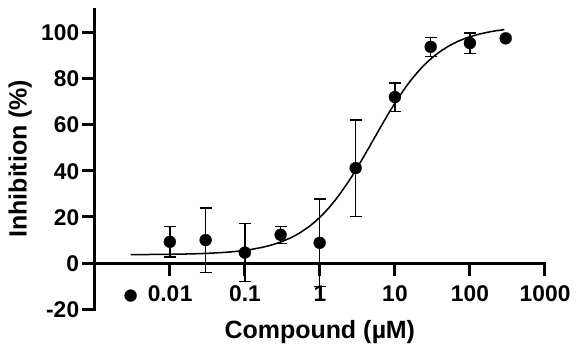 |  |
| 9 | **26** | **630** | 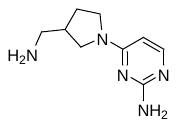 | 1.52 μM  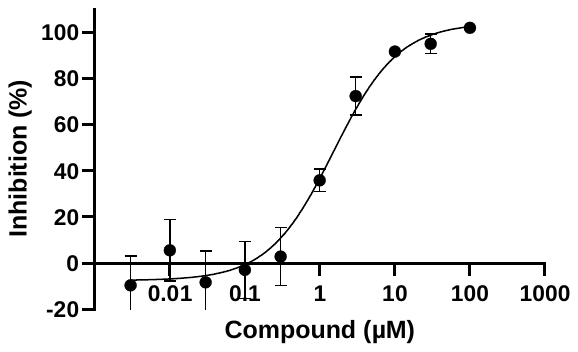 | 0.533 μM  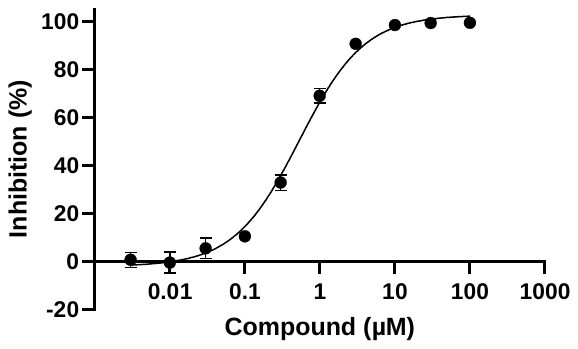 |
| 10 | **9** | **629** | 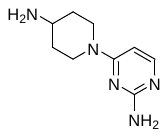 | 1.31 μM  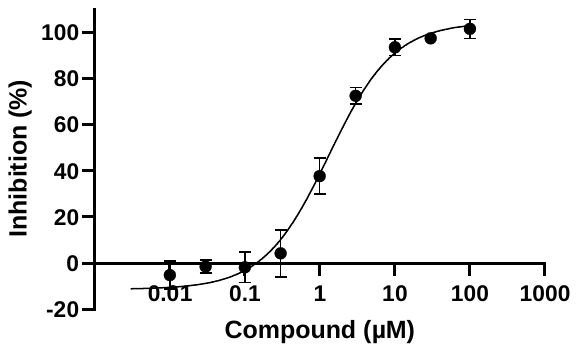 | 0.420 μM  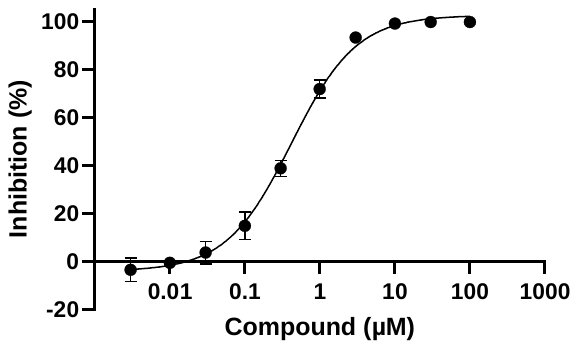 |
| 11 | **27** | **631** | 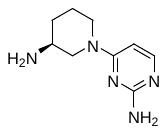 | 2.76 μM  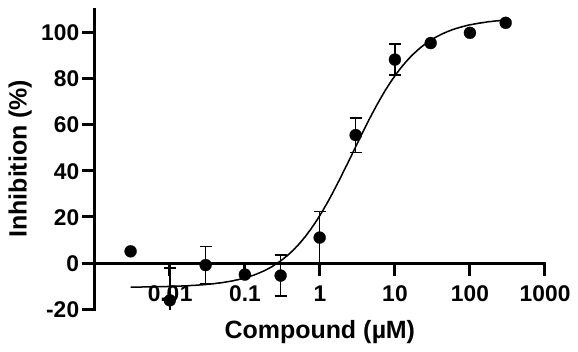 |  |
| 12 | **28** | **634** | 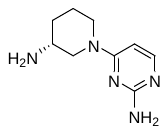 | 7.25 μM  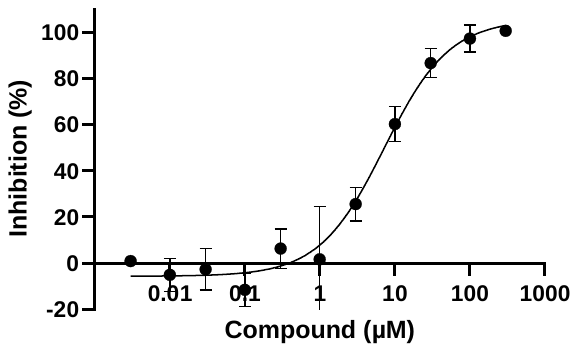 |  |
| 13 | **29** | **736** | 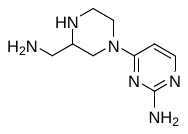 | 3.78 μM  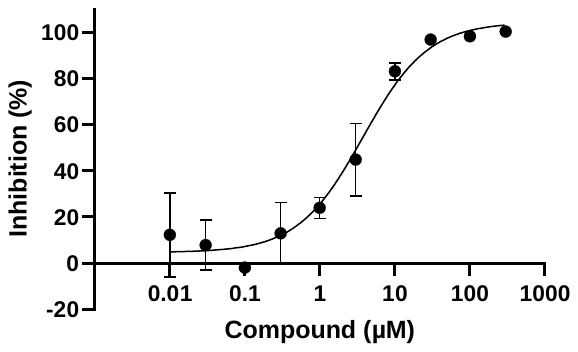 |  |
| 14 | **10** | **616** | 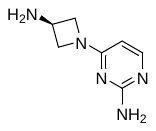 | 18.8 μM  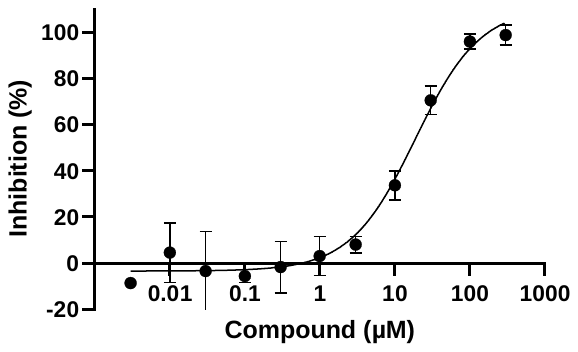 |  |
| 15 | **30** | **622** | 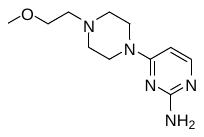 | >100 μM  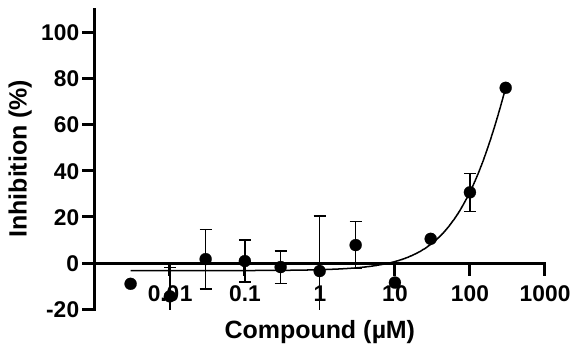 |  |
| 16 | **31** | **627** | 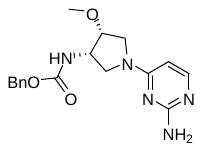 | >100 μM  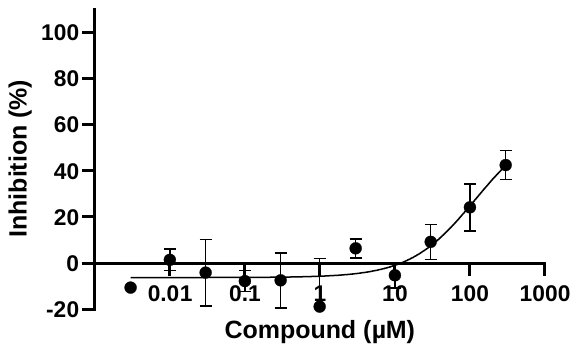 |  |
| 17 | **32** | **713** | 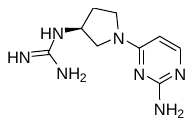 | 55.3 μM  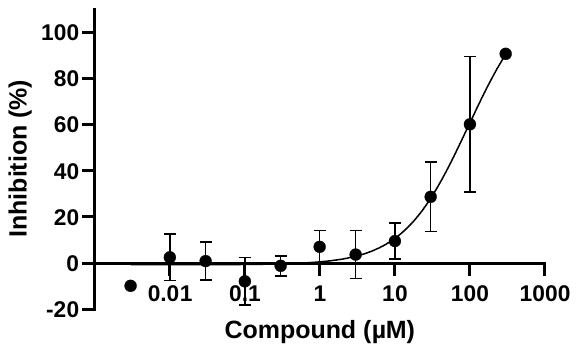 |  |
| 18 | **8** | **735** | 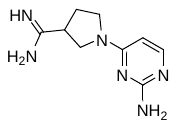 | 41.3 μM  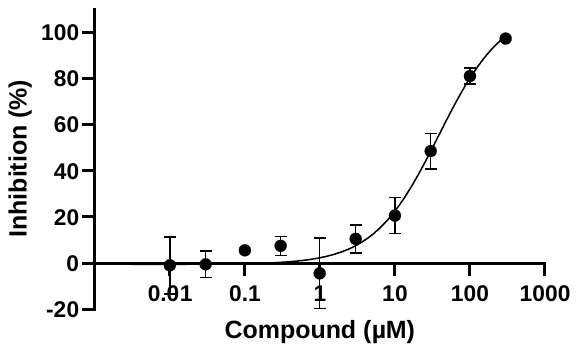 |  |
| 19 | **33** | **636** | 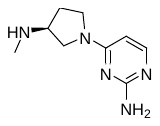 | 6.46 μM  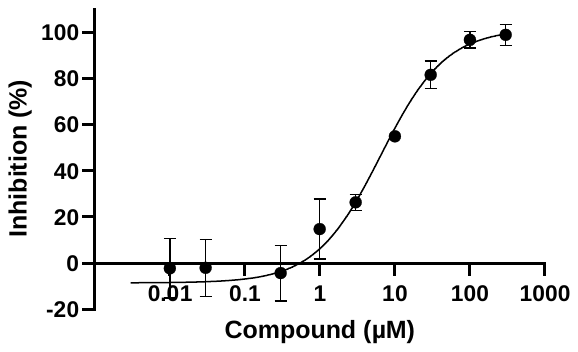 |  |
| 20 | **34** | **623** | 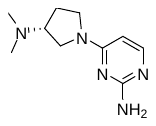 | 29.8 μM  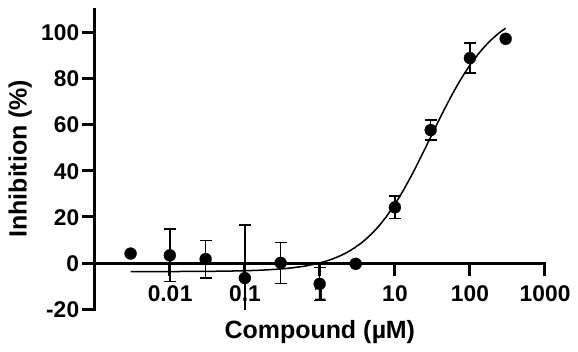 |  |
| 21 | **35** | **637** | 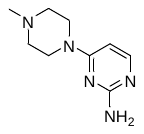 | 44.5 μM  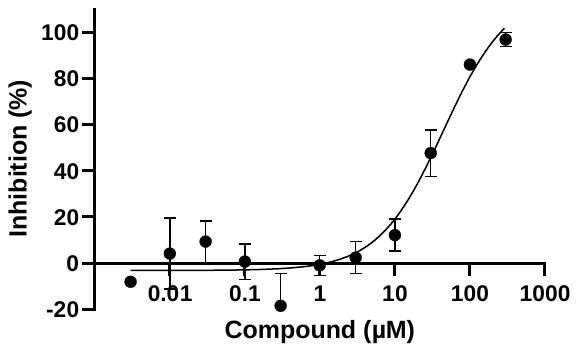 |  |
| 22 | **36** | **632** | 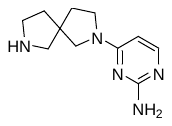 | 17.0 μM  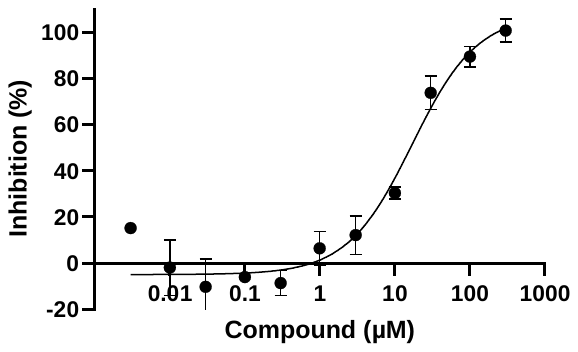 |  |
| 23 | **37** | **633** | 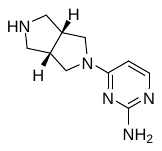 | 30.5 μM  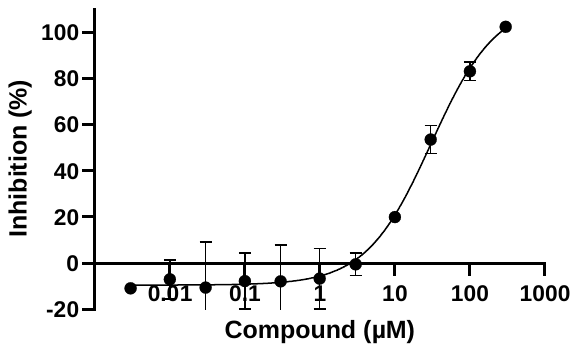 |  |
| 24 | **11** | **624** | 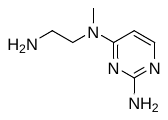 | 0.755 μM  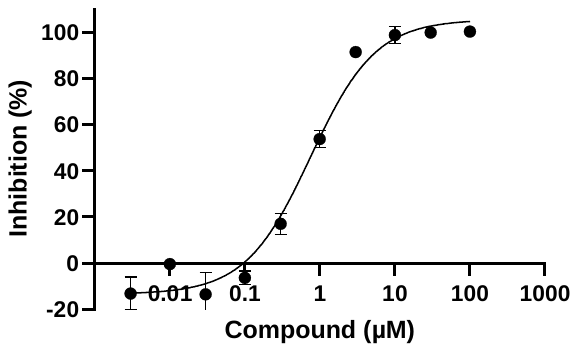 | 0.209 μM  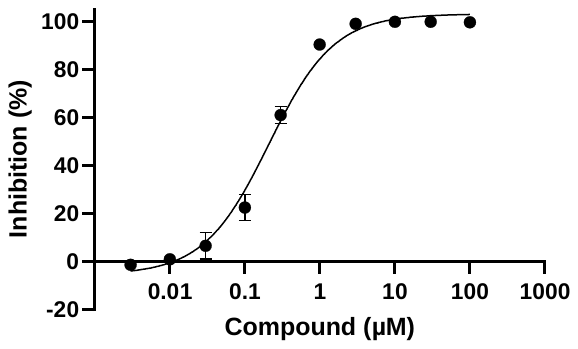 |
| 25 | **13** | **652** | 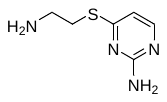 | Not active  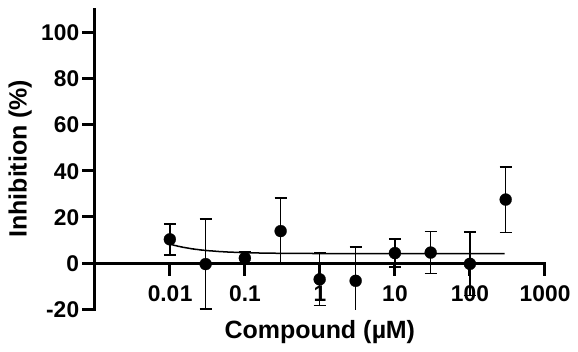 |  |
| 26 | **12** | **660** | 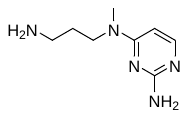 | 39.1 μM  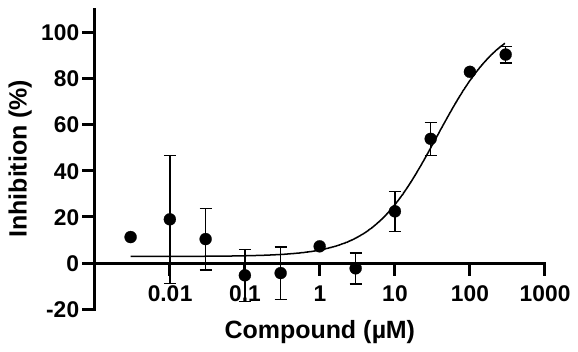 | 9.73 μM  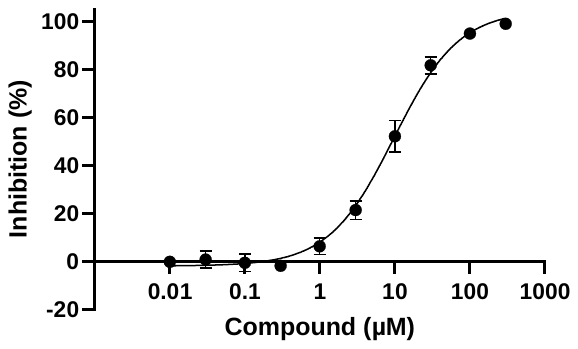 |
| 27 | **38** | **661** | 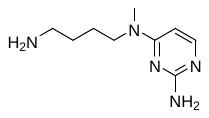 | 94.5 μM  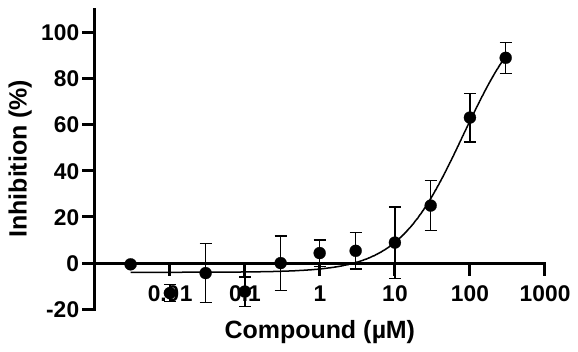 |  |
| 28 | **39** | **659** | 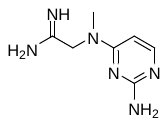 | Not active  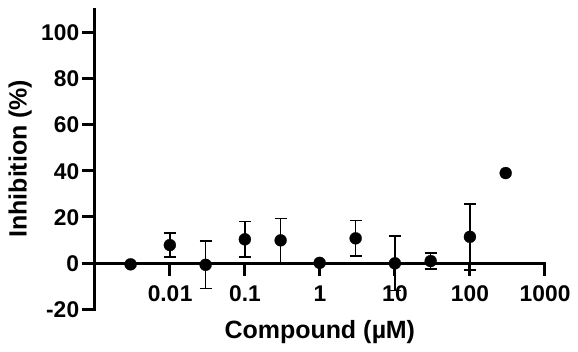 |  |
| 29 | **14** | **651** | 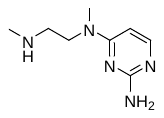 | 6.66 μM  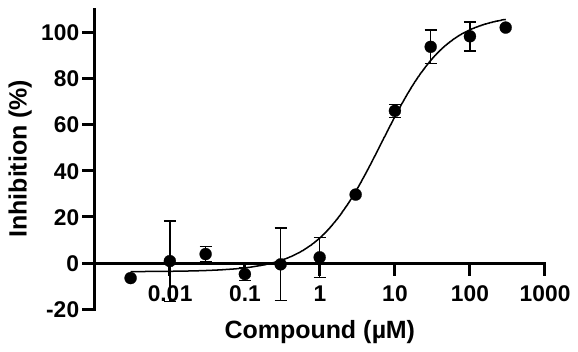 | 1.96 μM  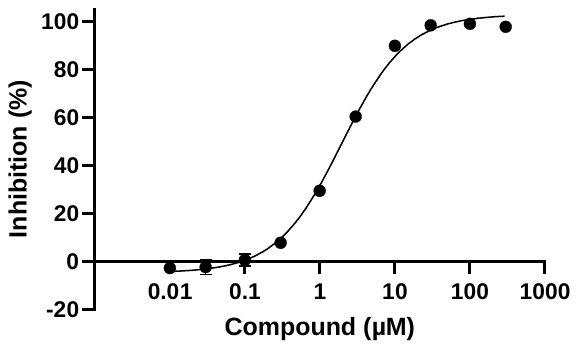 |
| 30 | **40** | **705** | 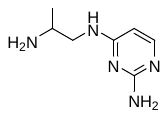 | 15.2 μM  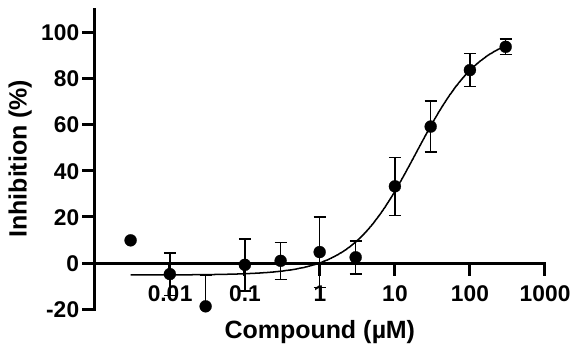 |  |
| 31 | **41** | **704** | 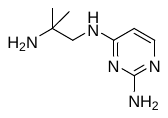 | >100 μM  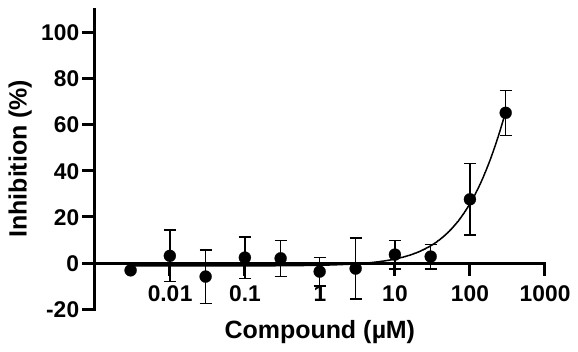 |  |
| 32 | **42** | **664** | 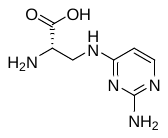 | 77.3 μM  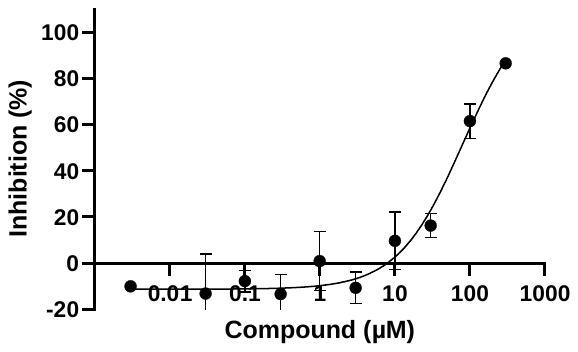 |  |
| 33 | **43** | **714** | 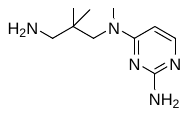 | >100 μM  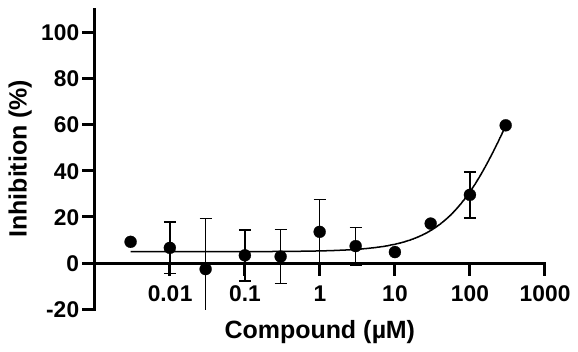 |  |
| 34 | **44** | **663** | 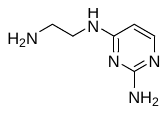 | 1.66 μM  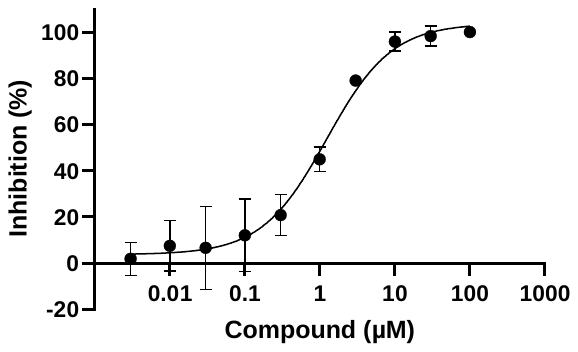 | 0.438 μM  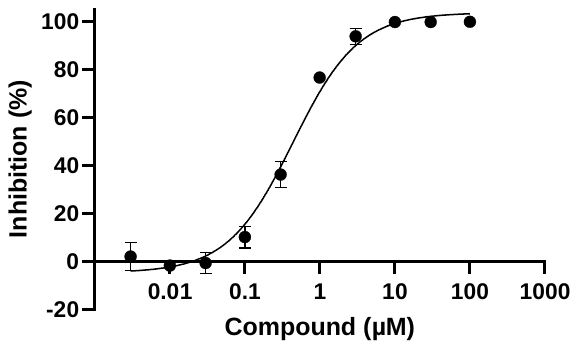 |
| 35 | **45** | **665** | 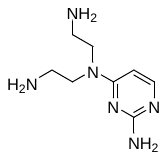 | 0.341 μM  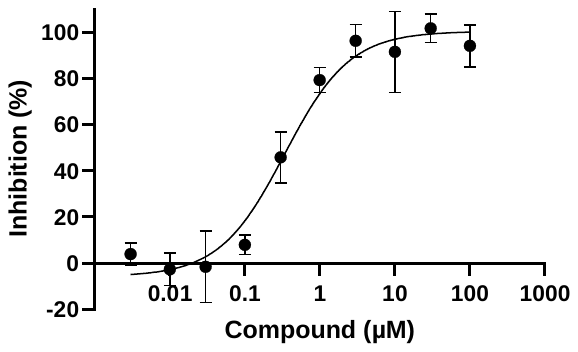 | 0.178 μM  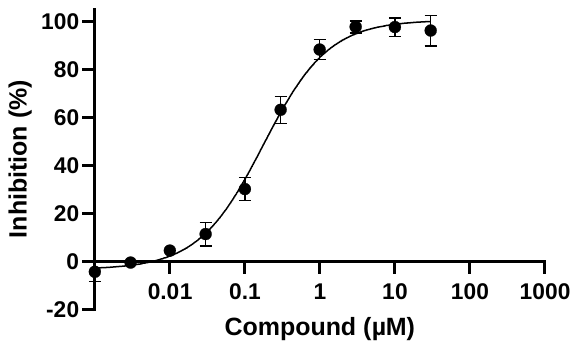 |
| 36 | **46** | **708** | 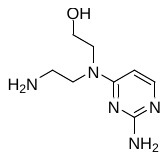 | 2.43 μM  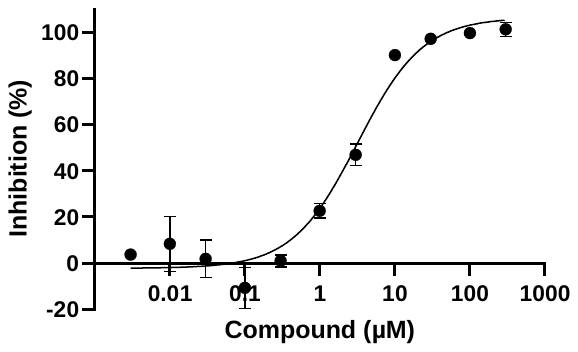 | 0.706 μM  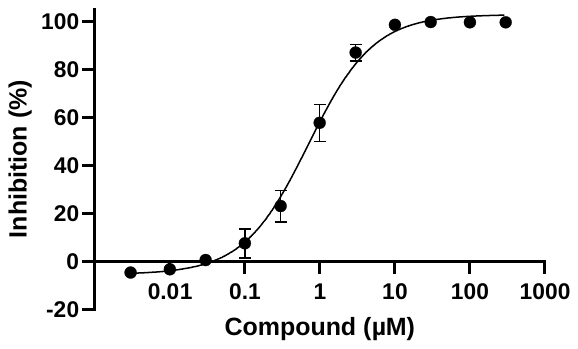 |
| 37 | **22** | **662** | 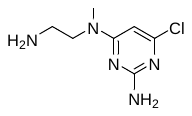 | Not active  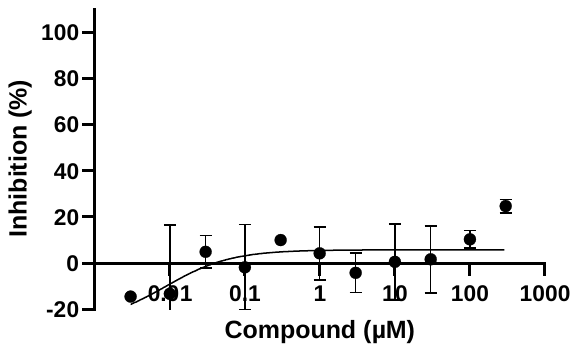 |  |
| 38 | **47** | **639** | 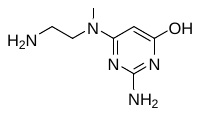 | >100 μM  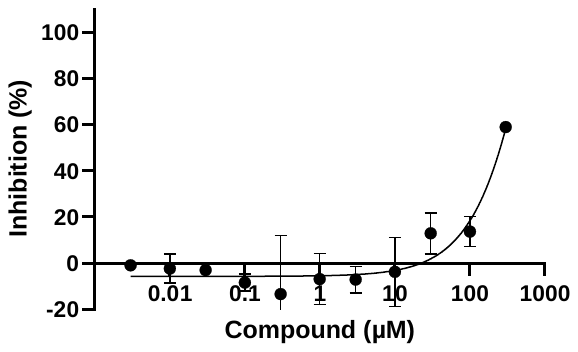 |  |
| 39 | **19** | **710** | 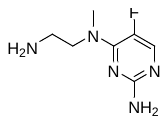 | Not active  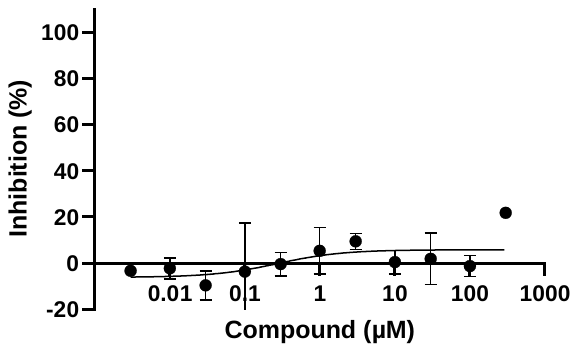 |  |
| 40 | **20** | **715** | 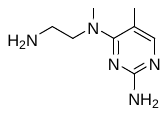 | 1.42 μM  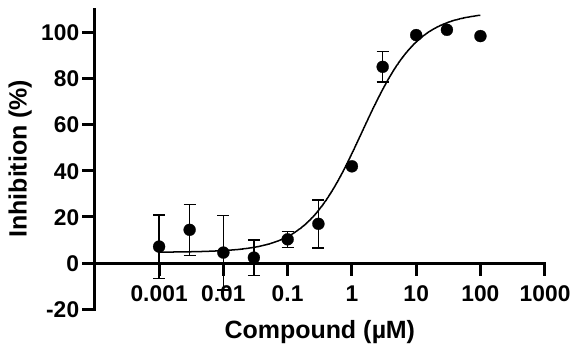 |  |
| 41 | **21** | **775** | 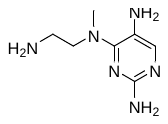 | 1.73 μM  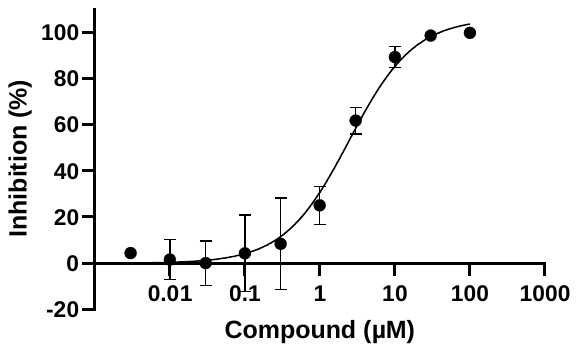 | 0.519 μM  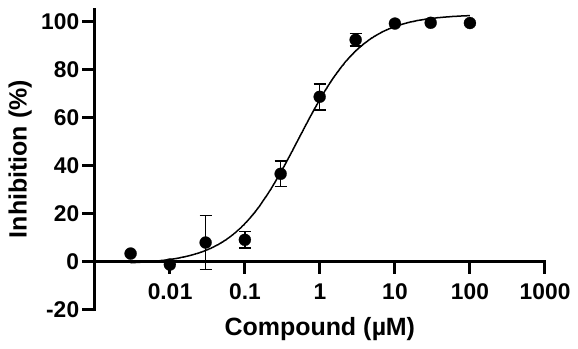 |
| 42 | **48** | **776** | 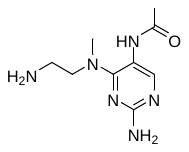 | >100 μM  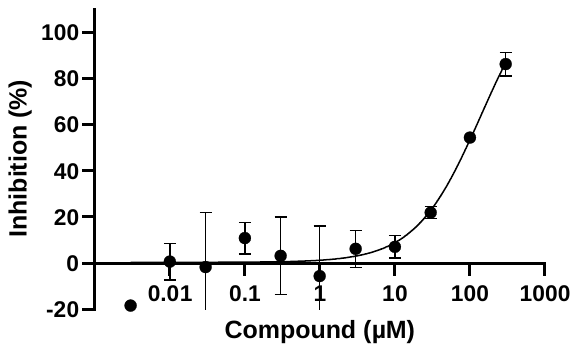 |  |
| 43 | **49** | **777** | 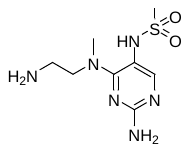 | >100 μM  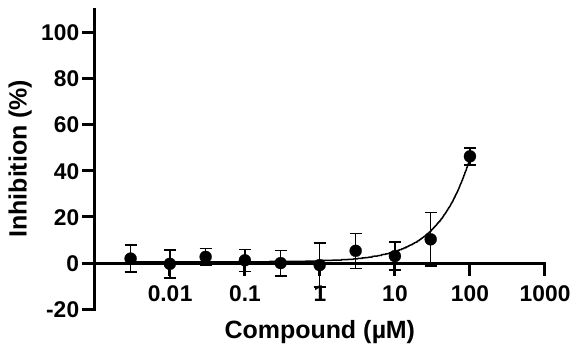 |  |
| 44 | **23** | **646** | 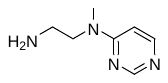 | >100 μM |  |
| 45 | **24** | **656** |  | >100 μM |  |
| 46 | **50** | **778** |  | >100 μM |  |
| 47 | **15** | **649** |  | >100 μM |  |
| 48 | **16** | **650** |  | >100 μM |  |
| 49 | **17** | **666** |  | 0.301 μM | 0.163 μM |
| 50 | **18** | **700** |  | 40.1 μM |  |
| 51 | **51** | **644** |  | >100 μM |  |
| 52 | **52** | **742** |  | 40.2 μM |  |
| 53 | **53** | **774** |  | 7.22 μM | 2.13 μM |
| 54 | **54** | **779** |  | 1.29 μM |  |
| 55 | **55** | **701** |  | Not active |  |
| 56 | **56** | **783** |  | 0.0433 μM | 0.0216 μM |

**Methods**

**Plasmid map**

Plasmid insert sequences for *in vitro* transcription of group I intron self-splicing constructs matched to *Candida albicans* SC5314 used in this study. C133T mutation was implemented from published sequence. The splicing kinetics between published transcript and transcript in this study were confirmed as identical.

| Plasmid | Encoded intron/exon | Insert sequence |
| --- | --- | --- |
| pJH102 | C.a.mtLSU | TAATACGACTCACTATAGGGTGATTTCTTCAGCATAGAGTGCCGATAACGGATCAAAGTTACGCTAGGGATAAACTGTTAGTCCCTTTATATAGAGATATATAGAATAACAATCGGGTGAATTGCAAGAATATCACCCACCTCCTTTGGAGGTGGGTGACAATTTGCAGCGAAGTATATATTAGCTTAATAATAAAATATTATTTAGATATATAAACGTTCAACGACTAGAAGGTGAGTAAGCTAACAATAACCCTTCCACGAGCGCCCGACATCTTAATGAACCAACAATTACAATTAATTAAAGGTCATTATGATGATGACATAGTCTGAACTAAATAGTGATATTTAGATATAATATTTAAATGATATTATGATAACAAAATTGAACAGGGTAATACCGCTAGAGAGTTGATATCGTCAGCGGTGTTTGCCACCGGATCC |

**Table S2**. Primer design for PCR and qPCR

| **Experiment** | **Name** | **Sequence** |
| --- | --- | --- |
| PCR | Total mtLSU (f) | GACCAAGTACAGTAAGACTAG |
|  | Total mtLSU (r) | CTTTGATCCGTTATCGGCAC |
|  | Unspliced mtLSU_lane2 (f) | GACCAAGTACAGTAAGACTAG |
|  | Unspliced mtLSU_lane2 (r) | CTTGCAATTCACCCGATTG |
|  | Unspliced mtLSU_lane3 (f) | GACCAAGTACAGTAAGACTAG |
|  | Unspliced mtLSU_lane3 (r) | GGTATTACCCTGTTATCCCTAG |
|  | Ligated exon (f) | GACCAAGTACAGTAAGACTAG |
|  | Ligated exon (r) | CGATATCAACTCTCTAGCGG |
|  | ACT1 (f) | GGAAGCTGCTGGTATTGA |
|  | ACT1 (r) | TAGACAATGAAGCCAAGATAGA |
|  | PGK1 (f) | CGCCAAAGATGGTGAAATC |
|  | PGK1 (r) | CTTCGAGACCAACCATAGAG |
| qPCR | Total RNA (f) | GACTTACAAGTCGACTTAGTACG |
|  | Total RNA (r) | CTTTGATCCGTTATCGGCAC |
|  | Unspliced RNA (f) | GACCAAGTACAGTAAGACTAG |
|  | Unspliced RNA (r) | TTCACCCGATTGTTATTCTATAT |
|  | ACT1 (f) | GGAAGCTGCTGGTATTGA |
|  | ACT1 (r) | TAGACAATGAAGCCAAGATAGA |

**Chemical Synthesis and Characterization**

**Materials and Methods**

All reactions were performed under a nitrogen atmosphere except for hydrogenations. Anhydrous solvents and 4M HCl in Dioxane were purchased from Sigma-Aldritch® (St. Louis, MS) and Fisher Scientific (Waltham, MA)®. Building blocks were obtained from either Combi-Blocks® (San Deigo, CA) or Ambeed® (Arlington Hts, IL). Palladium catalysts were purchased from TCI® USA. Additional reagents were sourced from TCI ®(Philadelphia, PA) and Oakwood Chemical® (Estill, SC). NMR Solvents were obtained from Cambridge Isotope Laboratories ® (Tewksbury, MA) and NMR spectra recorded on a Bruker® AdvanceCore 400 MHz spectrometer. HPLC analysis was performed on either an Agilent ® 1100 or 1260 system.

**Abbreviations Used**

ACN, acetonitrile

BOC, t*ert*-butoxycarbonyl

Cbz, benzyloxycarbonyl

CHCl_3_, chloroform

DBU, 1,8-Diazabicyclo(5.4.0)undec-7-ene

DCM, dichloromethane

DIPEA, N, N-Diisopropylethylamine

equiv., equivalents

EtOH, ethanol

HPLC, High-Performance Liquid Chromatography

IPA, isopropanol

M, molar

MeOH, methanol

MPLC, medium pressure chromatography

MsCl, methanesulfonyl chloride

NMR, Nuclear Magnetics Resonance

Pd/C, Palladium on carbon, 10% by weight

SnAR, Nucleophilic Aromatic Substitution

TEA, triethylamine

TFA, trifluoroacetic acid

TFAA, trifluoroacetic acid anhydride

THF, tetrahydrofuran

TLC, thin layer chromatography

vol., volumes

**General Procedures:** Many of the compounds described are synthesized using several common conditions. Synthetic procedures for compounds will refer to the general procedures used in their synthesis using the capital letters assigned to the steps below.

General Procedure A: SnAR of Heterocycles and Amines

To a glass pressure vessel equipped with stir bar was added the chloroheterocycle (1.0 equiv.) and the amine or derived HCl salt (1.0 equiv.). The reactants were suspended in an equal volume of EtOH and DIPEA to afford a 0.7 M solution. The pressure vessel was capped and heated at 100 °C for 16 hours with stirring. The reaction was then cooled to room temperature. If product precipitated the solid was filtered, rinsed with 2 reaction volumes of EtOH, and dried *in vacuo* to afford sufficiently pure product (>95% HPLC). If the product remained in solution the reaction solvent was removed *in vacuo* and the residue dissolved in 10 reaction volumes of DCM. The organic layer was washed with saturated aqueous NaHCO_3_ (3x 3 vol.), brine (1x 3 vol.), dried over anhydrous MgSO_4_, filtered, and the solvent was removed *in vacuo*. The residue was purified by MPLC using silica gel and a MeOH/DCM gradient.

HPLC conditions:

Column: Phenomenex Synergi: 4µm Polar-RP 80 Å 250 x 4.6 mm

Mobile Phase: A – water with 0.1% TFA, B – ACN with 0.1% TFA

Flow rate: 1.0 mL/minute

Gradient: 5-95% B from 0-10 minutes, 95% B from 10-13 minutes

General Procedure B: Cbz deprotection

A Cbz protected amine (1.0 equiv.) was placed in a round bottom flask with a stir bar and suspended in EtOH to afford a 0.165 M solution. The flask was purged with vacuum and backed filled 3x with N_2_(g) before the addition of Pd/C (0.1 equiv. 10% by weight) and Pd(OH)_2_/C (0.05 equiv. 20% by weight). A T-valve adapter equipped with a H_2_(g) balloon was fitted and the reaction was purged with vacuum and backed filled (3x) with H_2_(g) before stirring at 40 °C for 2.5 hours. After that time the reaction was cooled to room temperature and purged with vacuum before being backed filled 3x with N_2_(g). The reaction was filtered through a Celite ® pad, and the pad was rinsed with 3 reaction volumes of MeOH. The filtrate was concentrated *in vacuo*, dissolved in MeOH and treated with 4 M HCl in dioxanes (2.2 equiv.). The solvent was removed *in vacuo* to provide product as a hydrochloride salt.

General Procedure C: Boc deprotection

A Boc protected amine (1.0 equiv.) was placed in a round bottom flask with a stir bar and suspended in 4 M HCl in dioxanes to afford a 0.15 M solution. The mixture was heated to 40 °C for 2.5 hours before cooling to room temperature. The solid was filtered off and rinsed with one reaction volume of THF to furnish product as a hydrochloride salt.

General Procedure D: Nitro deprotection

A nitroheterocycle (1.0 equiv.) was added to a degassed solution of MeOH and Pd/C (0.16 equiv. 10% by weight) to afford a 0.078 M solution. A T-valve adapter equipped with a H_2_(g) balloon was fitted and the reaction was purged with vacuum and backed filled 3x with H_2_(g) before stirring at 23 °C for 3.5 hours. After that time the reaction was purged with vacuum before being backed filled 3x with N_2_(g). The reaction was filtered through a Celite ® pad, and the pad was rinsed with 3 reaction volumes of MeOH. The filtrate was concentrated *in vacuo* to provide product.

**4-(3-(aminomethyl)azetidin-1-yl)pyrimidin-2-amine dihydrochloride (25·2HCl):** This compound was synthesized using general procedures **A** and **C**, with yields of 59 and 94% respectively.

**^1^H NMR** (400 MHz, D_2_O) δ 7.54 (d, J = 7.3 Hz, 1H), 5.94 (d, J = 7.4 Hz, 1H), 4.44 – 4.29 (m, 2H), 4.06 – 3.92 (m, 2H), 3.31 (d, J = 7.6 Hz, 2H), 3.21 – 3.04 (m, 1H).

**^13^C NMR** (101 MHz, D_2_O) δ 161.54, 154.63, 140.62, 94.81, 53.38, 52.88, 41.97, 27.05.

**LCMS-ESI** m/z calculated for C_8_H_13_N_5_ [M+H^+^] 180.12, found 180.1.

**4-(3-(aminomethyl)pyrrolidin-1-yl)pyrimidin-2-amine dihydrochloride (26·2HCl):** This compound was synthesized using general procedures A and B, with yields of 60 and 77% respectively.

**^1^H NMR** (400 MHz, D_2_O) δ 7.72 (d, J = 6.8 Hz, 1H), 6.08 (d, J = 6.8 Hz, 1H), 3.96 – 3.59 (m, 2H), 3.53 (dt, J = 11.2, 8.0 Hz, 1H), 3.29 (dd, J = 11.2, 8.0 Hz, 1H), 3.23 – 3.09 (m, 2H), 2.78 – 2.64 (m, 1H), 2.44 –2.20 (m, 1H), 1.98 – 1.73 (m, 1H).

**^13^C NMR** (101 MHz, D_2_O) δ 160.47, 158.74, 148.43, 96.33, 49.69, 46.07, 41.45, 36.15, 28.19.

**LCMS-ESI** m/z calculated for C_9_H_15_N_5_ [M+H^+^] 194.14, found 194.2.

**(S)-4-(3-aminopiperidin-1-yl)pyrimidin-2-amine dihydrochloride (27·2HCl):** This compound was synthesized using general procedures A and B, with yields of 75 and 78% respectively.

**^1^H NMR** (400 MHz, D_2_O) δ 7.82 (d, *J* = 6.6 Hz, 1H), 6.27 (d, *J* = 6.6 Hz, 1H), 4.15 (dd, *J* = 13.4, 3.4 Hz, 1H), 3.80 (dt, *J* = 13.4, 4.6 Hz, 1H), 3.51 (dd, *J* = 13.4, 8.1 Hz, 1H), 3.46-3.33 (m, 2H), 2.21 – 2.10 (m, 1H), 1.93 – 1.61 (m, 3H).

**^13^C NMR** (101 MHz, D_2_O) δ 162.87, 161.34, 154.15, 95.02, 46.83, 46.73, 44.29, 28.37, 21.54.

**LCMS-ESI** m/z calculated for C_9_H_15_N_5_ [M+H^+^] 194.14, found 194.2.

**N^4^-(3-amino-2,2-dimethylpropyl)pyrimidine-2,4-diamine (29·2HCl):** This compound was synthesized using general procedures **A** and **D**, with yields of 76 and 82% respectively.

**^1^H NMR** (400 MHz, D_2_O) δ 7.82 (d, *J* = 6.7 Hz, 1H), 6.26 (d, *J* = 6.7 Hz, 1H), 4.11 – 4.03 (m, 1H), 3.20 (ddd, *J* = 13.6, 10.5, 3.3 Hz, 1H), 3.14 – 2.97 (m, 5H), 2.83 (ddd, *J* = 13.3, 10.4, 3.5 Hz, 1H).

**^13^C NMR** (101 MHz, D_2_O) δ 162.61, 160.85, 153.27, 95.02, 52.24, 46.24, 44.09, 42.99, 41.03.

**LCMS-ESI** m/z calculated for C_9_H_17_N_5_ [M+H^+^] 209.15, found 209.1.

**4-(4-(2-methoxyethyl)piperazin-1-yl)pyrimidin-2-amine dihydrochloride (30·2HCl):** This compound was synthesized using general procedure A in 26% yield.

**^1^H NMR** (400 MHz, DMSO) δ 7.73 (d, *J* = 6.0 Hz, 1H), 5.98 (d, *J* = 6.0 Hz, 1H), 5.93 (s, 2H), 3.49-3.40 (dt, 6H), 3.22 (s, 3H), 2.50 – 2.45 (m, 2H), 2.41 (t, *J* = 5.1 Hz, 4H).

**^13^C NMR** (101 MHz, DMSO) δ 162.95, 162.28, 156.76, 93.01, 69.87, 57.98, 57.01, 52.82, 43.20.

**LCMS-ESI** m/z calculated for C_11_H_19_N_5_O [M+H^+^] 238.17, found 238.2.

**(benzyl ((3S,4R)-1-(2-aminopyrimidin-4-yl)-4-methoxypyrrolidin-3-yl)carbamate (31):** This compound was synthesized using general procedure A in 76% yield.

**^1^H NMR** (400 MHz, DMSO) δ 7.72 (d, *J* = 5.9 Hz, 1H), 7.43 (d, *J* = 7.8 Hz, 1H), 7.38-7.34 (m, *J* = 4H), 7.34 – 7.29 (m, 1H), 5.95 (s, 2H), 5.70 (d, *J* = 5.8 Hz, 1H), 5.06 (s, 2H), 4.38-4.4.11 (s, 1H), 3.99-3.80 (s, 1H), 3.73 – 3.28 (m, 4H), 3.28 (s, 3H).

**^13^C NMR** (101 MHz, DMSO) δ 163.32, 161.11, 156.39, 156.26, 137.54, 128.82, 128.28, 128.21, 93.98, 78.58, 65.88, 57.29, 52.21, 48.93, 48.61.

**LCMS-ESI** m/z calculated for C_17_H_21_N_5_O_3_ [M+H^+^] 344.17, found 344.3.

**(S)-1-(1-(2-aminopyrimidin-4-yl)pyrrolidin-3-yl)guanidine dihydrochloride (32·2HCl):**

**^1^H NMR** (400 MHz, D_2_O) δ 7.69 (dd, *J* = 7.5, 4.0 Hz, 1H), 6.25 (dd, *J* = 20.1, 7.5 Hz, 1H), 4.43 – 4.31 (m, 1H), 4.04 – 3.55 (m, 4H), 2.52-2.30 (m,1H), 2.30 – 2.12 (m, 1H).

**^13^C NMR** (101 MHz, D_2_O) δ 160.61, 160.48, 156.38, 154.38, 140.81, 140.77, 96.92, 96.81, 52.08, 51.85, 50.97, 50.28, 45.23, 45.21, 30.35, 29.74. (all peaks doubled)

**LCMS-ESI** m/z calculated for C_9_H_15_N_7_ [M+H^+^] 222.15, found 222.2.

**(S)-4-(3-(methylamino)pyrrolidin-1-yl)pyrimidin-2-amine dihydrochloride (33·2HCl):** This compound was synthesized using general procedures **A** and **C**, with yields of 94 and 80% respectively.

**^1^H NMR** (400 MHz, D2O) δ 7.71 (d, J = 7.5 Hz, 1H), 6.26 (t, J = 7.4 Hz, 1H), 4.15 – 3.99 (m, 2H), 3.97 – 3.75 (m, 3H), 2.83 (d, J = 3.2 Hz, 3H), 2.68 – 2.48 (m, 1H), 2.44 – 2.25 (m, 1H).

**^13^C NMR** (101 MHz, D2O) δ 160.77, 160.65, 154.42, 141.22, 96.79, 96.73, 58.00, 57.34, 48.81, 48.65, 45.03, 44.96, 31.55, 31.47, 27.42, 26.79. (all peaks doubled except for 154.42 and 141.22).

**LCMS-ESI** m/z calculated for C9H15N5 [M+H+] 194.14, found 194.2

**(R)-4-(3-(dimethylamino)pyrrolidin-1-yl)pyrimidin-2-amine (34·2HCl):** This compound was synthesized using general procedure A in 45% yield.

**^1^H NMR** (400 MHz, MeOD) δ 7.71 (d, *J* = 6.2 Hz, 1H), 5.87 (d, *J* = 6.2 Hz, 1H), 4.01-3.51 (m, 2H), 3.43 – 3.33 (m, 1H), 3.23-3.13 (m, 1H), 2.94-2.79 (m, 1H), 2.33 (s, 6H), 2.30-2.19 (m, 1H), 1.94-1.77(s, 1H).

**^13^C NMR** (101 MHz, DMSO) δ 162.54, 160.37, 155.12, 93.64, 64.51, 50.07, 45.01, 43.76, 29.20.

**LCMS-ESI** m/z calculated for C_10_H_17_N_5_ [M+H^+^] 208.16, found 208.3.

**4-(4-methylpiperazin-1-yl)pyrimidin-2-amine (35):** This compound was synthesized using general procedure **A** in 73% yield.

**^1^H NMR** (400 MHz, DMSO) δ 7.74 (d, *J* = 6.0 Hz, 1H), 6.00 (d, *J* = 6.0 Hz, 1H), 5.95 (s, 2H), 3.49 (t, *J* = 5.2 Hz, 4H), 2.31 (t, *J* = 5.1 Hz, 4H), 2.19 (s, 3H).

**^13^C NMR** (101 MHz, DMSO) δ 162.93, 162.32, 156.74, 93.07, 54.29, 45.76, 43.10.

**LCMS-ESI** m/z calculated for C_9_H_15_N_5_ [M+H^+^] 194.14, found 194.3

**4-(2,7-diazaspiro[4.4]nonan-2-yl)pyrimidin-2-amine dihydrochloride (36·2HCl):** This compound was synthesized using general procedures A and C, with yields of 90 and 95% respectively.

**^1^H NMR** (400 MHz, D_2_O) δ 7.66 (dd, *J* = 7.5, 4.1 Hz, 1H), 6.22 (dd, *J* = 16.5, 7.5 Hz, 1H), 3.84 – 3.62 (m, 4H), 3.52 (d, *J* = 6.4 Hz, 2H), 3.40 (d, *J* = 6.7 Hz, 2H), 2.29 – 2.05 (m, 4H).

**^13^C NMR** (101 MHz, D_2_O) δ 160.44, 160.34, 154.42, 154.36, 146.92, 140.67, 140.62, 96.99, 96.77, 66.57, 55.22, 54.97, 52.30, 52.24, 48.28, 47.46, 46.47, 45.08, 33.90, 33.56, 33.47, 33.44. (All peaks doubled due to dynamics except 66.57)

**LCMS-ESI** m/z calculated for C_11_H_17_N_5_ [M+H^+^] 220.16, found 220.2.

**4-((3aR,6aS)-hexahydropyrrolo[3,4-c]pyrrol-2(1H)-yl)pyrimidin-2-amine dihydrochloride (37·2HCl):** This compound was synthesized using general procedures A and C, with yields of 62 and 100% respectively.

**^1^H NMR** (400 MHz, D_2_O) δ 7.68 (d, *J* = 7.5 Hz, 1H), 6.24 (d, *J* = 7.5 Hz, 1H), 4.01 – 3.88 (m, 2H), 3.82 – 3.60 (m, 4H), 3.44 – 3.24 (m, 4H).

**^13^C NMR** (101 MHz, D_2_O) δ 160.63, 154.40, 140.90, 97.00, 50.43, 50.22, 49.44, 41.29, 40.57. (Peaks at ~50.3 and 41.0 doubled)

**LCMS-ESI** m/z calculated for C_10_H_15_N_5_ [M+H^+^] 206.14, found 206.2.

**N^4^-(4-aminobutyl)-N4-methylpyrimidine-2,4-diamine (38·2HCl):** This compound was synthesized using general procedures **A** and **C**, with yields of 100 and 57% respectively.

**^1^H NMR** (400 MHz, D_2_O) δ 7.63 (dd, *J* = 7.6, 6.6 Hz, 1H), 6.36 (dd, *J* = 33.1, 7.7 Hz, 1H), 3.82 – 3.53 (m, 2H), 3.18 (d, *J* = 18.9 Hz, 2H), 3.06 (q, *J* = 7.2 Hz, 2H), 1.88 – 1.54 (m, 4H).

**^13^C NMR** (101 MHz, D_2_O) δ 162.25, 162.00, 154.32, 154.06, 140.75, 140.58, 95.88, 66.56, 50.10, 48.34, 39.19, 36.08, 35.79, 23.99, 23.86, 23.73, 23.19. (all peaks doubled except 95.88, 39.19)

**LCMS-ESI** m/z calculated for C_8_H_13_N_7_ [M+H^+^] 196.16, found 196.3

**2-((2-aminopyrimidin-4-yl)(methyl)amino)acetimidamide (39)**

**^1^H NMR** (400 MHz, MeOD) δ 7.87 (d, *J* = 6.3 Hz, 1H), 6.20 (d, *J* = 6.4 Hz, 1H), 4.56 (s, 2H), 3.15 (s, 3H).

**^13^C NMR** (101 MHz, DMSO) δ 167.93, 162.61, 162.11, 156.26, 93.52, 48.92, 36.43.

**LCMS-ESI** m/z calculated for C_7_H_12_N_6_ [M+H^+^] 181.12, found 181.1

**N^4^-(2-aminopropyl)pyrimidine-2,4-diamine dihydrochloride (40·2HCl):** This compound was synthesized using general procedures **A** and **C**, with yields of 96 and 44% respectively.

**^1^H NMR** (400 MHz, D_2_O) δ 7.61 (d, *J* = 7.3 Hz, 1H), 6.22 (d, *J* = 7.3 Hz, 1H), 3.84 – 3.63 (m, 3H), 1.37 (d, *J* = 6.5 Hz, 3H).

**^13^C NMR** (101 MHz, D_2_O) δ 164.43, 155.23, 140.58, 98.75, 47.64, 43.34, 15.35.

**LCMS-ESI** m/z calculated for C_7_H_13_N_5_ [M+H^+^] 168.12 found 168.1.

**N^4^-(2-amino-2-methylpropyl)pyrimidine-2,4-diamine dihydrochloride (41·2HCl):** This compound was synthesized using general procedures **A** and **C**, with yields of 95 and 74% respectively.

**^1^H NMR** (400 MHz, D_2_O) δ 7.63 (d, *J* = 7.3 Hz, 1H), 6.24 (d, *J* = 7.3 Hz, 1H), 3.73 (s, 2H), 1.44 (s, 6H).

**^13^C NMR** (101 MHz, D_2_O) δ 164.67, 155.23, 140.73, 98.72, 55.66, 47.69, 22.84.

**LCMS-ESI** m/z calculated for C_8_H_15_N_5_ [M+H^+^] 182.14 found 182.1.

**(S)-2-amino-3-((2-aminopyrimidin-4-yl)amino)propanoic acid dihydrochloride (42·2HCl):** This compound was synthesized using general procedures **A** and **C**, with yields of 33 and 59% respectively.

**^1^H NMR** (400 MHz, D_2_O) δ 7.63 (d, *J* = 7.5 Hz, 1H), 6.22 (d, *J* = 7.3 Hz, 1H), 4.28 (dd, *J* = 6.4, 3.9 Hz, 1H), 4.17 (dd, *J* = 15.0, 3.9 Hz, 1H), 3.93 (dd, *J* = 15.0, 6.4 Hz, 1H).

**^13^C NMR** (101 MHz, D_2_O) δ 170.36, 164.63, 155.19, 140.91, 98.66, 53.44, 40.25.

**LCMS-ESI** m/z calculated for C_7_H_11_N_5_O_2_ [M+H^+^] 198.10, found 198.1.

**N^4^-(2-aminoethyl)pyrimidine-2,4-diamine dihydrochloride (44·2HCl):** This compound was synthesized using general procedures **A** and **C**, with yields of 69 and 56% respectively.

**^1^H NMR** (400 MHz, D_2_O) δ 7.59 (d, J = 7.3 Hz, 1H), 6.20 (d, J = 7.3 Hz, 1H), 3.80 (t, J=5.8 Hz, 2H), 3.29 (t, J = 5.8 Hz, 2H).

**^13^C NMR** (101 MHz, D_2_O) δ 164.29, 155.22, 140.49, 98.78, 39.14, 37.95.

**LCMS-ESI** m/z calculated for C_6_H_11_N_5_ [M+H^+^] 154.11, found 154.1.

**N^4^,N^4^-bis(2-aminoethyl)pyrimidine-2,4-diamine trihydrochloride (45·3HCl):** This compound was synthesized using general procedures **A** and **C**, with yields of 77 and 96% respectively.

**^1^H NMR** (400 MHz, D_2_O) ^1^H NMR (400 MHz, D_2_O) δ 7.85 (d, *J* = 7.5 Hz, 1H), 6.51 (d, *J* = 7.6 Hz, 1H), 4.07 (t, *J* = 6.0 Hz, 2H), 3.95 (t, *J* = 7.3 Hz, 2H), 3.44-3.31 (m, 4H).

**^13^C NMR** (101 MHz, D_2_O) ^13^C NMR (101 MHz, D_2_O) δ 164.17, 154.43, 143.01, 95.46, 45.56, 45.46, 37.77, 36.19. (all aliphatic peaks doubled).

**LCMS-ESI** m/z calculated for C_8_H_16_N_6_ [M+H^+^] 197.15, found 197.2

**2-((2-aminoethyl)(2-aminopyrimidin-4-yl)amino)ethan-1-ol dihydrochloride (46·2HCl):** This compound was synthesized using general procedures **A** and **C**, with yields of 65 and 98% respectively.

**^1^H NMR** (400 MHz, MeOD) δ 7.66 (d, *J* = 6.3 Hz, 1H), 6.00 (d, *J* = 6.3 Hz, 1H), 3.69 (t, *J* = 5.7 Hz, 2H), 3.57 (d, *J* = 6.0 Hz, 4H), 2.86 (t, *J* = 6.6 Hz, 2H).

**^13^C NMR** (101 MHz, MeOD) δ 162.95, 162.51, 154.70, 93.74, 59.42, 50.66, 50.25, 38.94.

**LCMS-ESI** m/z calculated for C_8_H_15_N_5_O [M+Na^+^] 220.12 found 220.0.

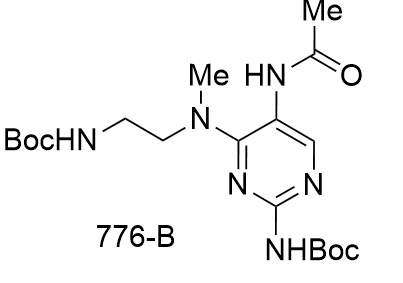

The Boc protected starting material was synthesized as detailed in (Cite first paper)

**tert-butyl N-{4-[(2-{[(tert-butoxy)carbonyl]amino}ethyl)(methyl)amino]-5-acetamidopyrimidin-2-yl}carbamate** (**48-B)**: tert-butyl N-{5-amino-4-[(2-{[(tert-butoxy)carbonyl]amino}ethyl)(methyl)amino]pyrimidin-2-yl}carbamate (1 eq., 75 mg, 0.2 mmol) was dissolved in DCM (0.94 mL), TEA (3 eq., 59.53 mg, 0.082 mL, 0.59 mmol)and Ac2O (3 eq., 60.058 mg, 0.055 mL, 0.59 mmol) were added. DMAP (0.2 eq., 4.79 mg, 0.039 mmol) was added and the reaction was stirred at rt for 1.5 hours.

After that time the reaction was diluted with DCM (10ml) and washed with NaHCO3 (3x 5 ml) and brine (1x 10 ml). The organic layer was dried over MgSO4, filtered, and concentrated. The residue was purifed on a 12 g biotage column using a gradient of 2->20 % MeOH in DCM over 10 col. vol. holding at 20% for an additional 2 col. vol. tert-butyl N-{4-[(2-{[(tert-butoxy)carbonyl]amino}ethyl)(methyl)amino]-5-acetamidopyrimidin-2-yl}carbamate (22.5 mg, 0.053 mmol, 27.029%) and tert-butyl N-[5-(N-acetylacetamido)-4-[(2-{[(tertbutoxy) carbonyl]amino}ethyl)(methyl)amino]pyrimidin-2-yl]carbamate (69 mg, 0.15 mmol, 75.42%) were isolated as white solids.

**N-{2-amino-4-[(2-aminoethyl)(methyl)amino]pyrimidin-5-yl}acetamide dihydrochloride (48·2HCl):** The final step was performed using general procedure **C**, the yield was 93%

**^1^H NMR** (400 MHz, D_2_O) δ 7.74 (s, 0H), 4.03 (t, *J* = 5.9 Hz, 1H), 3.38 (t, *J* = 5.8 Hz, 1H), 3.32 (s, 1H), 2.21 (s, 1H).

**^13^C NMR** (101 MHz, D_2_O) δ 175.53, 161.43, 152.74, 143.56, 107.79, 49.61, 37.29, 37.24, 21.70.

**LCMS-ESI** m/z calculated for C_9_H_16_N_6_O [M+H^+^] 225.15, found 225.0.

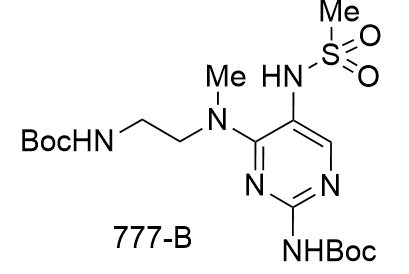

The Boc protected starting material was synthesized as detailed in (Cite first paper)

**tert-butyl N-{4-[(2-{[(tertbutoxy) carbonyl]amino}ethyl)(methyl)amino]-5-methanesulfonamidopyrimidin-2-yl}carbamate (49-B):** tert-butyl N-{5-amino-4-[(2-{[(tert-butoxy)carbonyl]amino}ethyl)(methyl)amino]pyrimidin-2-yl}carbamate (1 eq., 75mg, 0.2 mmol) was dissolved in 0.4 ml of DCM and pyridine (3 eq., 46.53 mg, 0.048 mL, 0.59 mmol). MsCl (1.2eq., 26.95 mg, 0.018 mL, 0.24 mmol) was dissolved in 0.55 ml of DCM . The reaction was stirred at RT for 3hours.

After that time the reaction was diluted with DCM (10ml) and washed with NaHCO3 (1x 5 ml), the organic layer was dried over MgSO4, filtered, and concentrated. The reaction was purified on a 12g biotage column with a gradient of 2->20% MeOH in DCM over 12 column volumes. tert-butyl N-{4-[(2-{[(tertbutoxy) carbonyl]amino}ethyl)(methyl)amino]-5-methanesulfonamidopyrimidin-2-yl}carbamate (62.8 mg, 0.14 mmol, 69.54%) was obtained as a yellow solid.

**N-{2-amino-4-[(2-aminoethyl)(methyl)amino]pyrimidin-5-yl}methanesulfonamide dihydrochloride (49·2HCl):** The final step was performed using general procedure **C**, the yield was 35%

**^1^H NMR** (400 MHz, D_2_O) δ 7.95 (s, 1H), 4.06 (t, *J* = 6.0 Hz, 2H), 3.51 (s, 3H), 3.40 (m, 2 t, *J* = 6.0 Hz H), 3.23 (s, 3H).

**^13^C NMR** (101 MHz, D_2_O) δ 162.18, 152.69, 144.58, 107.01, 49.67, 39.05, 38.88, 37.11.

**LCMS-ESI** m/z calculated for C_8_H_16_N_6_O_2_S [M+H^+^] 261.11, found 261.2.

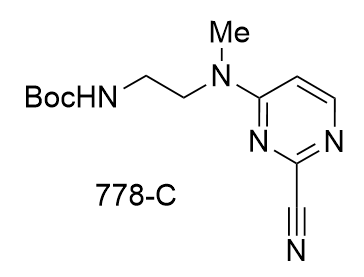

**tert-butyl N-{2-[(2-cyanopyrimidin-4-yl)(methyl)amino]ethyl}carbamate (50-C)** This step was performed using general procedure **A**, the yield was 88%

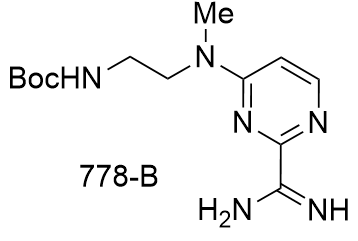

**N-{2-[(2-carbamimidoylpyrimidin-4-yl)(methyl)amino]ethyl}carbamate** **(50-B):** A solution of tert-butyl N-{2-[(2-cyanopyrimidin-4-yl)(methyl)amino]ethyl}carbamate (1 eq., 100 mg, 0.36 mmol) in MeOH (2.2 mL) was treated with 25% in MeOH NaOMe (2.5 eq., 194.802 mg, 0.901 mmol) and stirred at 50 °C (step: 1) for 2 hours (step: 1). NH4Cl (2.5 eq., 48.22 mg, 0.901 mmol) was added and the solution stirred at 65 °C (step: 2) for 2 hours (step: 2). The reaction was checked by TLC (20% MeOH/EtOAc or DCM, UV) which showed trace amounts of S1, intermediate, and P1 (baseline spot) present. The solvent was removed in vacuo and the crude residue (~200 mg) was dissolved in minimal 5% MeOH/EtOAc (flask rinsed with a little 10% MeOH/EtOAc) and purified by flash chromatography (12g silica, 5-20% MeOH/EtOAc 8CV, 20% 4CV, 25% MeOH/DCM 13CV) to provide tert-butyl N-{2-[(2-carbamimidoylpyrimidin-4-yl)(methyl)amino]ethyl}carbamate (68 mg, 0.23 mmol, 64.066%) as a pink solid.

**4-[(2-aminoethyl)(methyl)amino]pyrimidine-2-carboximidamide dihydrochloride (50·2HCl)** The final step was performed using general procedure **C**, the yield was 82%

**^1^H NMR** (400 MHz, D_2_O) δ 8.40 (d, *J* = 6.3 Hz, 1H), 6.95 (d, *J* = 6.4 Hz, 1H), 4.16-4.02 (m, 2H), 3.36 (t, *J* = 6.1 Hz, 2H), 3.19 (s, 3H).

**^13^C NMR** (101 MHz, D_2_O) δ 162.40, 160.14, 155.00, 151.41, 106.43, 46.02, 37.22, 35.63.

**LCMS-ESI** m/z calculated for C_8_H_14_N_6_ [M+H^+^] 195.14, found 195.2.

**N^6^-(2-aminoethyl)-N^6^-methyl-9H-purine-2,6-diaminedihydrochloride (51·2HCl):** This compound was synthesized using general procedures **A** and **C**, with yields of 68 and 100% respectively.

**^1^H NMR** (400 MHz, D_2_O) δ 8.12 (s, 1H), 4.41-3.91 (m, 2H), 3.53 (s, 3H), 3.39 (t, *J* = 5.7 Hz, 2H).

**^13^C NMR** (101 MHz, D_2_O) δ 154.48, 152.18, 146.52, 141.40, 99.99, 48.05, 37.64, 37.45.

**LCMS-ESI** m/z calculated for C_8_H_13_N_7_ [M+H^+^] 208.13, found 208.3

**tert-butyl (2-(2-amino-4-chloro-7H-pyrrolo[2,3-d]pyrimidin-7-yl)ethyl)carbamate (52-A):** To a room temperature solution of 4-Chloro-7H-pyrrolo[2,3-d]pyrimidine-2-amine ( 1.0 equiv, 500 mg, 2.97 mmol) in ACN (14.8 ml) was added tert-butyl N-(2-bromoethyl)carbamate (1.1 equiv., 731 mg, 3.26 mmol). Cs_2_CO_3_ (1.5 equiv., 1449 mg, 4.45 mmol) and NaI (1.2 equiv., 533 mg, 3.56 mmol) were added before the reaction was stirred for 16 hours at 75°C. After that time the reaction was cooled to room temperature, diluted with EtOAc (75 ml), washed with water (3x 20 ml), and brine (1x 20 ml). The organic layer was dried over MgSO_4_, filter, and the solvent removed *in vacuo*. The residue was purified was purified by flash chromatography (silica gel, 12-80% EtOAc/Hexanes) to provide tert-butyl (2-(2-amino-4-chloro-7H-pyrrolo[2,3-d]pyrimidin-7-yl)ethyl)carbamate ( 174 mg, 0.56 mmol, 9.0 %) as a white solid.

**tert-butyl (2-(2-amino-7H-pyrrolo[2,3-d]pyrimidin-7-yl)ethyl)carbamate (52-B):** (2-(2-amino-4-chloro-7H-pyrrolo[2,3-d]pyrimidin-7-yl)ethyl)carbamate (1.0 equiv., 173 mg, 0.55 mmol) was suspended in MeOH (5.5 ml ) and TEA (1.2 equiv. 67 mg, 93 μL 0.67 mmol) was added. The mixture was purged and back filled with N_2_ (3x) before the addition of Pd/C (0.1 equiv., 59 mg, 0.055 mmol). A t-valve adaptor with a H_2_ balloon was affixed and the reaction was purged and back filled with H_2_ (3x) . The reaction was stirred in a 40°C oil bath for 2 hours. After that time the reaction was cooled to room temperature, filter over a pad of Celite ®, and rinsed with MeOH. The solvent was removed *in vacuo* and the residue was purified by was purified was purified by flash chromatography (silica gel, 2-13% MeOH/DCM) to provide tert-butyl (2-(2-amino-7H-pyrrolo[2,3-d]pyrimidin-7-yl)ethyl)carbamate (117, mg, 0.42 mmol, 76%) as a white solid.

**7-(2-aminoethyl)-7H-pyrrolo[2,3-d]pyrimidin-2-amine (52·2HCl):** The final step was performed using general procedure **C**, the yield was 91%

**^1^H NMR** (400 MHz, D_2_O) δ 8.55 (s, 1H), 7.42 (d, J = 3.9 Hz, 1H), 6.72 (d, J = 3.9 Hz, 1H), 4.51 (t, J = 5.8 Hz, 2H), 3.53 (t, J = 5.8 Hz, 2H).

**^13^C NMR** (101 MHz, D_2_O) δ 155.46, 152.24, 138.72, 132.59, 112.54, 103.09, 41.96, 39.01.

**LCMS-ESI** m/z calculated for C_8_H_11_N_5_ [M+H^+^] 178.11, found 178.0.

**tert-butyl (2-(2-amino-4,4a,5,6-tetrahydro-7H-pyrrolo[2,3-d]pyrimidin-7-yl)ethyl)carbamate(53-A):** ter**t-**butyl (2-(2-amino-7H-pyrrolo[2,3-d]pyrimidin-7-yl)ethyl)carbamate (742-B) (1.0 equiv., 148 mg, 0.534 mmol) was dissolved in AcOH (0.5 ml) and EtOAc (0.5 ml). The mixture was purged and back filled with N_2_ (3x) before the addition of Pd/C, 10 wt% (0.2 equiv., 114 mg, 0.107 mmol). A t-valve adaptor with a H_2_ balloon was affixed and the reaction was purged and back filled with H_2_ (3x). The reaction was stirred vigorously at 40 °C for16 hours. After that time the reaction was filtered of Celite ®, the pad rinsed with MeOH, and the solvent removed *in vacuo*. 7-(2-aminoethyl)-4a,5,6,7-tetrahydro-4H-pyrrolo[2,3-d]pyrimidin-2-amine ( 150 mg ideal, 0.534, 100%) was isolated as a slightly overweight white solid. This material was used in the subsequent step without purification.

**7-(2-aminoethyl)-4a,5,6,7-tetrahydro-4H-pyrrolo[2,3-d]pyrimidin-2-amine (53·2HCl):** The final step was performed using general procedure **C**, the yield was 86%

**^1^H NMR** ^1^H NMR (400 MHz, D_2_O) δ 4.00 – 3.71 (m, 5H), 3.43 – 3.35 (m, 2H), 3.29 – 3.16 (m, 2H), 2.54 – 2.40 (m, 1H), 2.03 – 1.89 (m, 1H).

**^13^C NMR** ^13^C NMR (101 MHz, DMSO) δ 173.91, 160.76, 51.40, 42.47, 41.54, 36.41, 36.34, 24.91.

**LCMS-ESI** m/z calculated for C_8_H_15_N_5_ [M+H^+^] 182.14, found 182.0

**N^5^-(2-aminoethyl)-N^5^-methylpyridine-2,5-diamine dihydrochloride (55·3HCl):** This compound was synthesized using general procedures **D** and **A**, with yields of 88 and 96% respectively.

**^1^H NMR** (400 MHz, D_2_O) δ 7.81 (dd, *J* = 9.7, 2.9 Hz, 1H), 7.31 (d, *J* = 2.9 Hz, 1H), 7.04 (d, *J* = 9.7 Hz, 1H), 3.50 (t, *J* = 6.3 Hz, 2H), 3.25 (t, *J* = 6.3 Hz, 2H), 2.87 (s, 3H).

**^13^C NMR** (101 MHz, D_2_O) δ 148.58, 138.55, 136.23, 118.61, 114.65, 50.09, 38.33, 36.51.

**LCMS-ESI** m/z calculated for C_8_H_14_N_4_ [M+H^+^] 167.13, found 167.2

**S_N_Ar:** To a pressure tube was added 4-chloro-2-nitropyridine (790 mg, 5 mmol), di-tert-butyl (azanediylbis(ethane-2,1-diyl))dicarbamate (1.5 g, 5 mmol, 1.0 equiv.), DMSO (5 mL), and DIPEA (5 mL). Then the reaction was stirred at 100 ^o^C overnight. After cooling to room temperature, the reaction mixture was dissolved in EtOAc (50 mL). Then the reaction mixture was washed with saturated brine (50 mL X 3), dried over MgSO_4_, and filtrated. After evaporation, the residue was purified by silica gel chromatography and then PTLC (DCM: MeOH=10:1) to give product as yellow solid (1.1 g, 52%).

**Reduction:** The above nitrocyclic compound (1.0 equiv.) was added to a degassed solution of MeOH and Pd/C (0.16 equiv. 10% by weight) to afford a 0.078 M solution. A T-valve adapter equipped with a H_2_(g) balloon was fitted and the reaction was purged with vacuum quickly and backed filled 3x with H_2_(g) before stirring at 23 °C for 3.5 hours. After that time, the reaction was filtered through a Celite ® pad, and the pad was rinsed with 3 reaction volumes of MeOH. The filtrate was concentrated in vacuo to provide product without further purification (note: the product was unstable in silica gel, so LCMS was used to monitor the reaction).

**Boc Deprotection:** 1) A Boc protected amine (1.0 equiv.) was placed in a round bottom flask with a stir bar and suspended in 4 M HCl in dioxanes to afford a 0.15 M solution. The mixture was heated to 40 °C for 2.5 hours before cooling to room temperature. The solid was filtered off and rinsed with Et_2_O to furnish product as a hydrochloride salt.

2) A Boc protected amine (1.0 equiv.) was dissolved in DCM to afford a 0.15 M solution, followed by the addition of 3 equiv. TFA. The mixture was stirred at rt overnight. After removing the solvent, the resulting residue was washed with Et_2_O to furnish product as a TFA salt.

***N*^4^,*N*^4^-bis(2-aminoethyl)pyridine-2,4-diamine (56·2HCl):** This compound was synthesized using general procedures **A**, **D**, and **C**, with yields of 52%, 100%, and 94% respectively.

**HCl salt:** ^1^H NMR (400 MHz, D_2_O) δ 7.63 (d, *J* = 7.6 Hz, 1H), 6.51 (dd, *J* = 7.7, 2.7 Hz, 1H), 6.07 (d, *J* = 2.7 Hz, 1H), 3.86 (t, *J* = 7.2 Hz, 4H), 3.30 (t, *J* = 7.2 Hz, 4H). ^13^C NMR (151 MHz, D_2_O) δ 156.78, 153.49, 135.90, 100.70, 89.32, 46.92, 35.73.

**TFA salt:** ^1^H NMR (400 MHz, D_2_O) δ 7.55 (d, *J* = 7.6 Hz, 1H), 6.44 (dd, *J* = 7.6, 2.7 Hz, 1H), 5.99 (d, *J* = 2.6 Hz, 1H), 3.80 (t, *J* = 7.2 Hz, 4H), 3.24 (t, *J* = 7.1 Hz, 4H). ^13^C NMR (101 MHz, D_2_O) δ 156.68, 153.48, 135.80, 100.53, 89.18, 46.84, 35.63.

**HRMS-ESI** m/z calculated for C_9_H_18_N_5_^+^ [M+H]^+^ 195.1484, found 196.1558.
